# Neuronal Dot1l is a broad mitochondrial gene-repressor associated with human brain aging via H3K79 hypermethylation

**DOI:** 10.1101/2021.10.11.463907

**Authors:** H.J Van Heesbeen, L Von Oerthel, P.M De Vries, M.R.J Wagemans, M.P. Smidt

## Abstract

Methylation of histone 3 at lysine 79 (H3K79) and its catalyst, disrupter of telomeric silencing (Dot1l), have been coupled to multiple forms of stress like bioenergetic and ER challenges. However, studies on H3K79 methylation and Dot1l in the aging brain and neurons are very limited. This together with increasing evidence of a dynamic neuroepigenome made us wonder if H3K79 methylation and Dot1l could play unknown roles in brain aging and associated disorders. In aged humans, we found strong and consistent hypermethylation of H3K79 in neurons that accumulate lipofuscine, while neuronal Dot1l transcript abundance reacts to bioenergenic and oxidative challenges. Indeed, in dopaminergic neurons we found rapid global H3K79me turnover (<12h). While shortly after reduction of H3K79 methylation, synaptic transcripts decreased while mitochondrial genes, particularly respiratory chain transcripts increased. Strikingly, 6 months after reduction of Dot1l levels, almost solely a variety of mitochondrial genes linked to aging and Parkinsons disease remained increased. These profiles are in much detail inverse to those described in hallmark PD and aging studies and associate Dot1l and H3K79me with neuronal stress in the aging brain while putting Dot1l forward as dynamic master regulator of mitochondrial transcription in dopamine neurons.

## Introduction

The dynamic character of the epigenome in post-mitotic neurons has gained much appreciation the last decade, from a mere debate, to evidence for complete and ongoing replacement of histone subunits in post-mitotic neurons, regardless of their epigenomic code (1, 2). Epigenomic profiles in neurons may partly reflect a steady-state rather than a long-lasting rigid code per se. Such a dynamic balance of histone modification and eviction could shift in aging or diseased neurons, leading to global deregulation or facilitating homeostatic adaptation of neurons, influencing healthy lifespan and disease progression (3). In Parkinson’s Disease (PD), especially several mechanisms related to balanced histone acetylation and hydroxymethylation of DNA have been related to PD prevalence and disease mechanisms (4). However, related epigenetic mechanisms have been scarcely investigated, and many not at all. Here we have focused on methylation of lysine 79 on histone 3 (H3K79) which is exclusively targeted by Dot1l in mammalian cells (5). Dot1l is recruited towards (active) genes via direct binding to Polymerase II and towards acetylated histones through recruitment by the transcription factor Af9 (6, 7). Levels of Dot1l expression and recruitment have been related to an increase in methylation valence of H3K79 (8–10), generally leading to the accumulation of H3K79 dimethylation (H3K79me2). In an array of tested cell lines, H3K79me2 was found ubiquitously present whereas H3K79me3 is limited (11).

Dot1l/DOT1l has been coupled to Wnt signaling via binding to beta-catenin-containing complexes (12, 13). However, during the neuronal maturation phase in the mouse cortex Dot1l is strongly down-regulated (14). In addition, several studies have coupled Dot1l chromatin regulation to Sirt1 and other Sir(t) proteins, linking Dot1(l) mechanistically to key factors of aging and inflammatory pathways (15–17)

In cellular models, basal DOT1l levels and H3K79 methylation increase following cellular stress like responses to viral infections, by manipulation of bioenergetics (18–20), or by matrix hyaluronan, which depletion reduces DA cell loss and α-synuclein in a mouse model of synucleinopathy (21, 22). More recently, researching early life stress mouse models, both Dot1l and H3K79 methylation have been found to be long-lasting deregulated in medial spinal neurons (23). Finally, in neural stem cells Dot1l has been related to the ER stress response as well (24). Taken together, DOT1l and H3K79me2 levels generally increase following a broad range of cellular stressors.

Among the diverse neuronal types especially DA neurons located the substantia nigra pars compacta (SNc) are the most vulnerable to degenerate in PD, containing numerous arborizations that require structurally and bioenergetic support(25–27). Even though mechanistically and physiologically Dot1l and H3K79 methylation relate to key factors in aging and DA neurodegeneration, little is known about the roles and regulation of Dot1l and H3K79 methylation in (DA) neurons. Here, we have investigated H3K79 methylation levels in the aged and diseased human midbrain and DA neurons and studied the gene-regulatory roles of Dot1l in DA neurons of mice in depth.

We have found a highly consequent accumulation of H3K79 methylation in lipofuscine loaded neurons in the aged human midbrain, while H3K79 methylation showed a high turnover in mouse DA neurons. Furthermore, neuronal Dot1l transcript abundance increases both upon AMPK activation and oxidative stress, while transcriptome profiling of DA neurons found Dot1l as master regulator of mitochondrial transcription, especially repressing respiratory chain genes in DA neurons at physiological levels. Strikingly, in adult neurons mono-allelic deletion of Dot1l had remarkably influence on a set off genes related to mitochondria, aging and neurodegenerative diseases.

## Results

### H3K79me2 hypermethylation associates with lipofuscin in the aged human midbrain

Lipofuscine (LF) and neuromelanin (NM) containing macro-vacuole structures have been proposed as markers for vulnerable DA neurons in the midbrain (28). In addition, the formation and maturation of Lewy Bodies can involve stages of lysosomal dysfunction associating with lipofuscine accumulation (29). Densely folded autofluorescent protein aggregates in LF accumulate in lipid lysosomal spheres mostly the endoplasmic reticulum, emitting wavelengths of both 488nm and 555nm (30). As such, LF can be visualized as spherical yellow structures that surround nuclei (**Fig. 1A**). Since LF does not overlap with nuclei (**Fig. 1A**), we were able to investigate levels of histone modifications using immunological staining.

**Figure 1.**
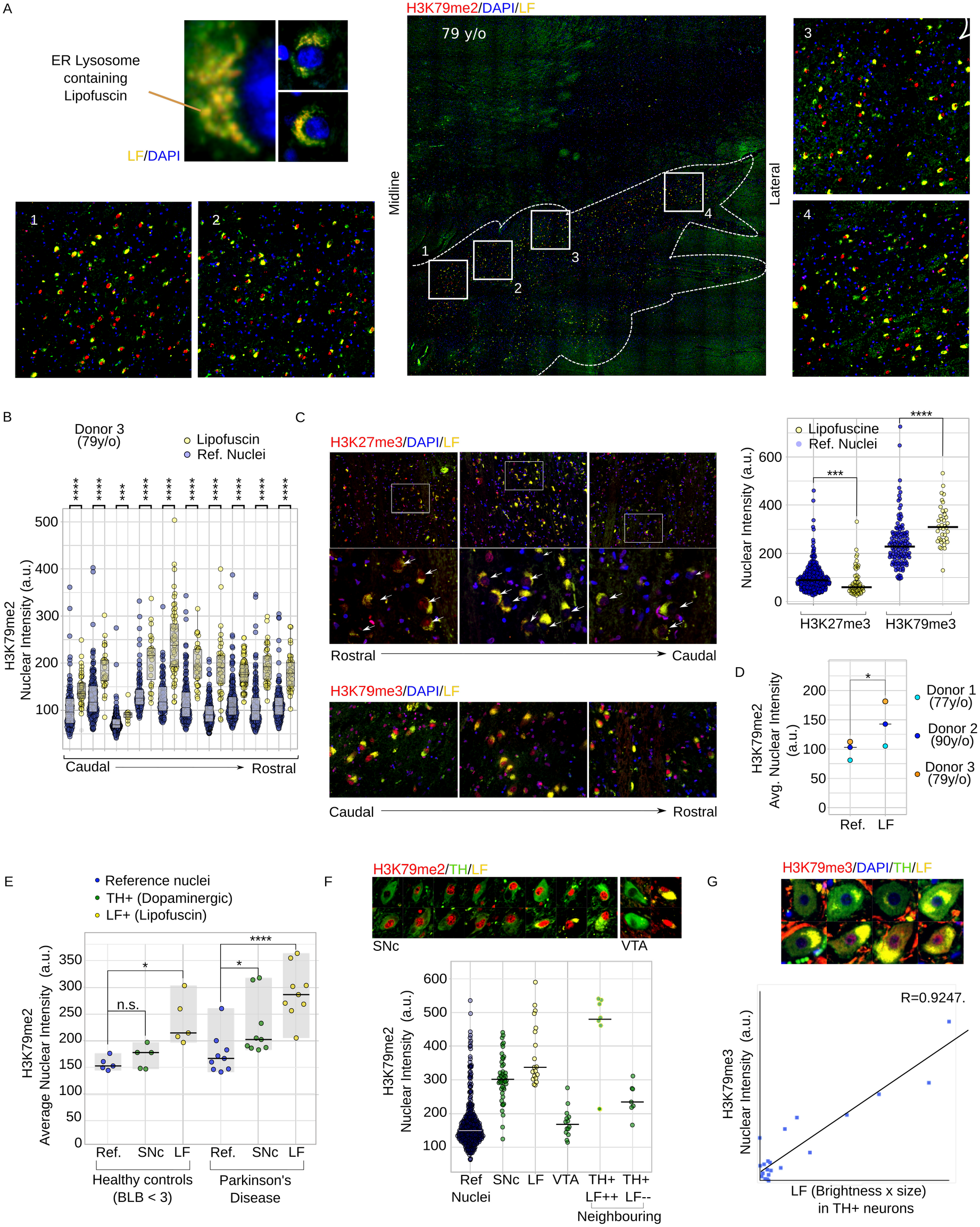
Increased H3K79me2 associates with Lipofuscin in the human midbrain. (**A**) Fluorescent image of typical red+green (yellow) autofluorescent lipid spheres containing accumulated dense protein (waste) structures in an ER-like morphology surrounding large nuclei (blue). Lipofuscin accumulation in a midbrain area overlapping with the SNc of a 79-year-old male with only minor loss of pigmented neurons, no Lewy-body pathology (**Table 1**) and no clinical PD but low TH levels (almost absent visible TH staining). White lines indicate the area with major LF accumulation. (**A1-4**) Magnification boxes in numbered 1-4 representing regions from medial to lateral showing the correlation between LF-autofluorescent neurons and H3K79me2 nuclear staining. Nuclear staining of DAPI (blue), H3K79me2 (red), and in LF (autofluorescence green/red-Yellow). (**B**) Quantification of sections from rostral to caudal. Each comparison shows LF associated nuclei and automatically selected surrounding reference nuclei from a single merged image. Mann-Whitney statistical tests were performed to compare the median of LF associated nuclear H3K79me2 staining with those of the surrounding reference nuclei for each photo. (**C**) Selection of images from a rostral to caudal series showing H3K79me3 (bottom), but not H3K27me3 (top), correlating to nuclei surrounded by LF. The upper right diagram shows quantification of H3K27me3 and H3K79me3 of adjacent coupes in LF and surrounding reference nuclei. Mann-Whitney statistical tests were used for median (of average per nuclei) levels of H3K79me3 and H3K27me3 surrounding nuclei in adjacent coupes. (**D**) Comparison between average H3K79me2 levels of reference and LF-associated nuclei measured from combined rostral and caudal midbrain sections of three aged human donors. A paired Students T-test was used to tests if average nuclear H3K79me2 levels are increased as compared to surrounding reference nuclei in three different donors. (**E**) Comparison of dopaminergic (TH+) and surrounding LF+ in non-Parkinson and PD human donors in de substantia nigra pars compact (SNc), compared to surrounding reference nuclei. Paired Student t-test were used for statistics. (**F**) Further dissecting cell types of one individual with high levels of H3K79me2 staining in TH+ SNc neurons but not those of the ventral tegmental area (VTA), functioning as an extra internal control. The bottom diagram shows the individual nuclear average H3K79me2 levels of reference nuclei (Ref. Nuclei), TH+ neurons in the substantia nigra (SNc), surrounding LF+/TH low/absent (LF (SNc)), VTA neurons surrounding the red nucleus (VTA (RN)) and neighboring neurons that show at least minimal TH staining but then in the mere absence (TH+ LF--) or strong presence (TH+/LF++) of LF. (**G**) Correlation (pearson) between the [size+intensity] of LF and nuclear H3K79me3 staining. Other statistics: * p=0.05-0.01, ** p=0.01-0.001, *** p=0.001-0.0001, **** p<0.0001. For (**E**), a post-hoc Bonferoni-correction for multiple testing was used.

In a 79-year-old male donor without pathological and cognitive signs of PD, we found a large number of LF surrounded nuclei stretching from the midline to lateral that associated with elevated H3K79me2 staining (**Fig. 1A**). Nuclear quantification revealed a comparable pattern from rostral to caudal of elevated H3K79me2. (**Fig. 1B**). We also investigated levels of the polycomb complex 2 histone mark H3K27me3. The reduction of H3K27me3 has been linked to senescent states and the upregulation of neurodegeneration-associated genes (31, 32). In contrary to H3K79me2, we have found that average nuclear H3K27me3 staining was lower in nuclei of LF containing cells as compared to surrounding controls while H3K79me3 was increased in sections adjacent to those used for H3K79me2 identification (**Fig. 1C, Sup. Fig. 1A/B**), together suggesting that H3K79 hypermethylation, but not generally H3 hypermethylation correlates to LF.

To investigate if the correlation between LF and H3K79 hypermethylation is a general phenomenon, we expanded our study with a 77 and 90-year-old donor confirming a consequent increase in H3K79me2 in the rostral and caudal midbrain in all investigated sections (**Fig. 1D, Sup. Fig. 1C/D**). This, regardless of the varying and scattered localization of LF+ cells. Overall, we only found remarkably few exceptions of LF+ cells showing less excessive H3K79me2 (**Sup. Fig. 1C/D**). In summary, LF-surrounded nuclei, including typical decondensed neuronal nuclei, have elevated levels of H3K79 methylation in aged individuals.

### H3K79me2 hypermethylation in TH+ DA neurons of PD patients is not generally increased in the absence of LF

Next, we investigated DA neurons and LF+ cells in the PD substantia nigra compacta (SNc). Again we found, without exception, H3K79 hypermethylation correlating with LF, both in healthy and PD individuals (**Fig. 1E, Sup. Fig. 2**). We also found an average increase in DA neurons from PD patients. However, the effect was much less profound (**Fig. 1E**). We did not find increased H3K79me2 in DA neurons in healthy individuals in the SNc, nor increased H3K79me3 in TH+ neurons without disease hallmarks (**Fig. 1E, Sup. Fig. S1E**). An overview of the levels in individual nuclei per donor can be found in **Supplemental Figure S2**.

We further investigated one PD donor with high levels of H3K79me2 in TH+ neurons in more depth by comparing the SNc DA neurons with those in the dorsal VTA that contain high levels of TH (**Sup. Fig 3**). We did not observe H3K79-hypermethylation in TH+, even though neighboring VTA LF+ cells and TH+ cells that contained LF showed similar patterns of a general increase relating to LF (**Fig. 1F**). Finally, we want to point out that TH/LF double-positive cells were scarce, especially in the ventral tier of the SNc, even though we could find them in abundance in neighboring cells. In rhesus monkeys, the VTA contains more LF+TH+ neurons while accumulation of aging markers inversely correlates with TH (33). Indeed, we did find several scattered sites on the border of the SNc/VTA, more ventral to the Red Nucleus, where reasonable numbers of LF+ DA neurons were found in proximity. There, H3K79me3 (next to H3K79me2) correlated with the size and intensity of the LF containing vacuoles (**Fig. 1G**). In those cells TH+ cells, levels of H3K79me3 only seemed to correlate to relatively high LF levels as compared to H3K79me2 in the previous datasets and generally in LF containing cells/neurons (**Fig. 1G**).

In summary, in all 17 human brain samples and in each section that we have investigated we have found H3K79me2 levels increased in cells that accumulated typical Lysosomal LF in ER-like patterns. Only in two individuals (both PD) had substantially elevated levels of H3K79me2 in TH+ DA neurons in the absence of LF, while LF and TH-staining was scarce, but in those neurons, a similar correlations was found.

### Neuronal Dot1l transcript abundance increases through H2O2 and AMPK activation

Several studies have suggested that cellular and mitochondrial stress may precede the upregulation of Dot1l as pointed out in the introduction, we wondered if these mechanisms are active in neurons too. To assess if bioenergetic manipulation or oxidative stress influences neuronal Dot1l transcript abundance, we have treated primary neurons both with the AMP mimicking AMPK activator AICAR and H2O2. After 6 hours Dot1l mRNA was significantly up-regulated in primary neurons following AICAR administration, while after 24 hours, both H2O2 and AICAIR showed increased Dot1l mRNA (**Fig. 2A**).

**Figure 2.**
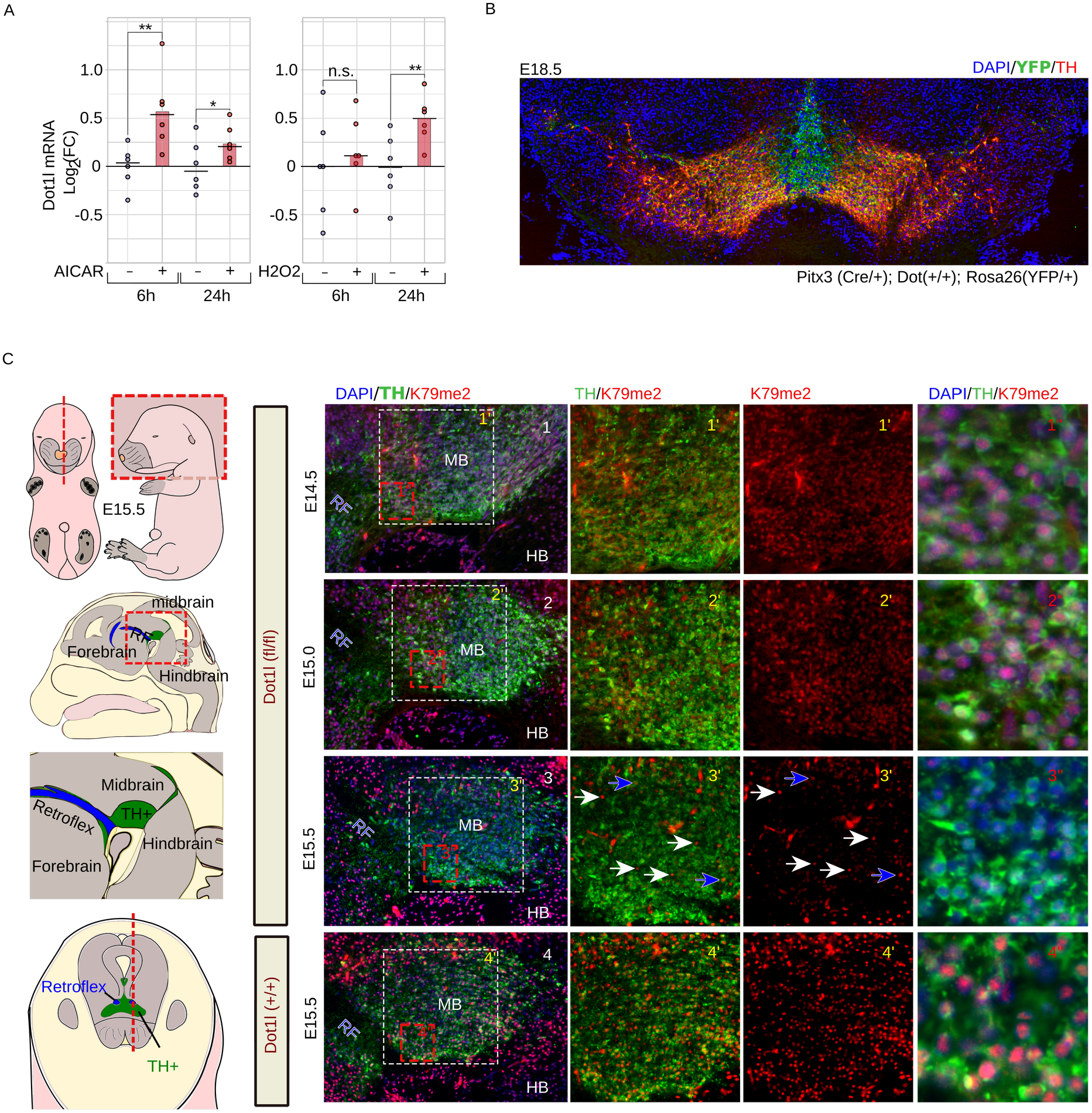
Neuronal *Dot1l* transcriptional regulation and H3K79me2 turnover. (**A**) Quantitative PCR of Dot1l mRNA levels in primary neurons either 6 or 24 hours following AICAR or H2O2 treatment with a final concentration of resp. 100 μM and 10μM. starting at DIV11. (**B**) Color combined image of a coronal midbrain section of an E18.5 *Pitx3*(*Cre*/+); *Dot1l*(+/+); *Rosa26*(*YFP*/+) embryo, stained for TH (Red), YFP (green) and DAPI (blue). (**C**) In the left panel a schematic representation of the location of the (RF), hindbrain (HB), and the DA (TH+) area at E15.5. In the right panel a time-line of sagital midbrain (MB) sections of *Pitx3*(*Cre*/+);*Dot1l*(*fl*/*fl*) at E14.5 (**C1**), E15.0(**C2**) and E15.5(**C3**) and *Dot1l*(+/+) at E15.5(**C4**) as control. All sections are matched by the presence of the retroflex (RF) and the location of the midbrain is marked MB. The first column (**C1-C4)** represents color combined overviews of the whole midbrain with DAPI (blue), Tyrosine Hydroxylase (TH, green), and H3K79me2 (red). Columns 2 and 3 (**C1’-C4’)** are magnifications that represent TH/H3K79me2 (left) and solely H3K79me2 (right) staining. Column 4 (**C1”-C4”**) shows overlays at higher magnifications presenting the nuclear depletion of H3K79me2 at E15.5 specifically in E15.5 *Pitx3*(*Cre*/+);*Dot1l*(*fl*/*fl*) embryos **(C3’’**). Blue arrows indicate the scarce TH+ neurons at E15.5 that still posses clear H3K79me2 staining, white arrows indicating non-Th+ cells. These cells contain clearly visible H3K79me2 staining in the middle of low-H3K79me2 stained Th+ neurons.

### Fast turnover of H3K79 methylation in post-mitotic DA neurons

To further investigate Dot1l function and H3K79 dynamics in post-mitotic DA neurons, we crossed mice that had loxp-sites flanking exon 5 of *Dot1l* (34) with mice that start expressing Cre-recombinase from the Pitx3 locus in post-mitotic DA precursors (35) while activating a reporter allele (Rosa26-lox-stop-lox-YFP, **Fig. 2 B**).

Even though Cre is active around E12.0 (35), we did not observe reduced K79 methylation at E15.0 in *Dot1l* floxed neurons (**Fig. 2C**). However, between E15.0 and E15.5 global H3K79 dimethylation rapidly declined in the majority of dopamine (DA) neurons (**Fig. 2C**). Only most rostro-lateral (minority), and in the most medial sections, we did find cells that were still H3K79me2 positive (**Sup. Fig. 4A/B**). We expect those to be the youngest neurons that may still have sufficient Dot1l levels. Similar results were obtained for H3K79me1 at E15.5 (**Sup. Fig. 4C/D**), together suggesting a rapid global turnover of H3K79 methylation. At later embryonic stages (E18.5), both me1 and me2 (not shown) remained absent in TH+ neurons, with only very sporadically a K79me1-positive DA neuron left (**Sup. Fig. 1E**).

### Complete Dot1l deletion leads to a rostro-lateral non-progressive developmental cell defect

We were uncertain if (complete) post-mitotic Dot1l deletion would alter the DA cell number/survival that could influence transcriptome profiling. Therefore, we have mapped expression levels and patterns of several key DA and subset markers. With qPCR on dissected midbrains, we found that transcript levels of critical DA genes were especially affected in full conditional Dot1l knock-outs (cKO) (*Th*, *Vmat2*, *Dat*, *Rspo2*, *Nurr1*, *Pitx3*, with *Girk2* and *Ahd2* following a trend), while heterozygous flox-mice were only minimally affected (**Fig. 3A**). In cKO mice, we also found the density of Th+ cells reduced in the compacted region of the substantia nigra, already at P7 (**Fig. 3B,C**). *Th* and *Ahd2 in situ hybridisation* (ISH) analyses of 6 months old mice showed a similar pattern with an *Ahd2-* positive DA neuronal population missing as well as a reduction of Th area/intensity in *Dot1l*(*fl*/*fl*) animals (**Fig. 3D**). Finally, we compared levels of mRNA between p40 and one year old *Dot1l*(*fl*/*fl*) mice but found no progressive loss of (or increase) of *Pitx3*, *Th*, *Adh2* or, resp. *Cck*. Together, we reasoned that postnatally, especially the heterozygous floxed mice were suited to further investigate the roles of Dot1l in modulating the DA transcriptome.

**Figure 3.**
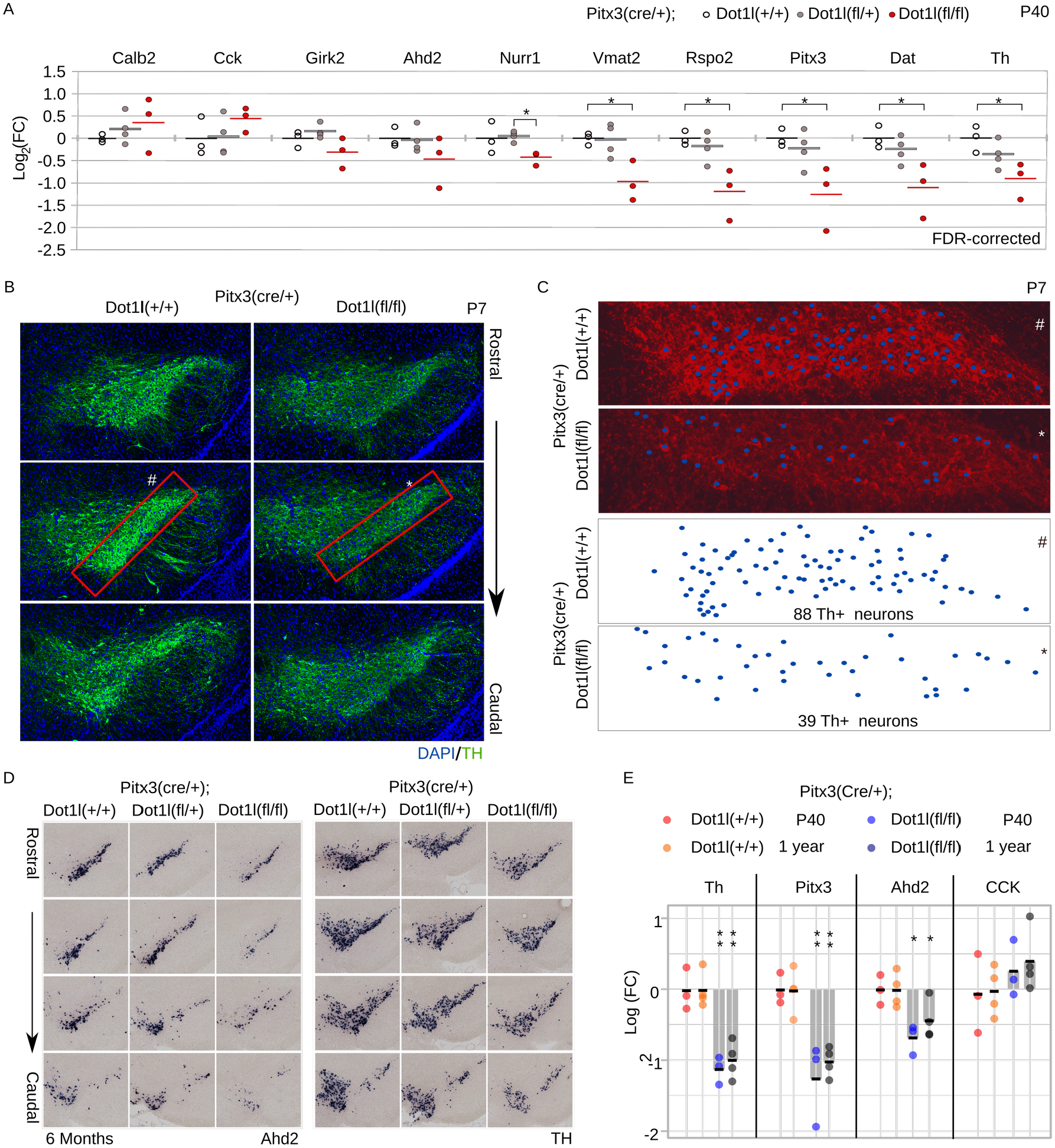
Neuronal defect in *Pitx3*(*Cre*/+);*Dot1l*(*fl*/*fl*) mice. (**A**) Quantitative PCR of dopaminergic and midbrain (subset) markers on dissected P40 midbrains. White dots represent *Dot1l*(+/+) (n=3), gray dots represent *Dot1l*(*fl*/+) (n=4), red dots represent *Dot1l*(*fl*/*fl*) (n=3), all in a *Pitx3*(*Cre*/+) background. Levels were normalized against *Tbp* transcript levels. (**B**) Immunohistochemistry for Th in P7 coronal sections of rostral *Pitx3*(*Cre*/+); *Dot1l*(+/+) and *Dot1l*(*fl*/*fl*) mice. (**C**) Mono color representation of marked sections (Red boxes) in ‘B’, with ‘#’, indicating the densely TH populated region of the *Pitx3*(*Cre*/+); *Dot1l*(+/+) SNc and with ‘*’ the *Dot1l*(*fl*/*fl*). The blue dots are a graphical representation of counted TH-+ neurons that suggest that not only a level, but also a decrease in cell number occurs. (**D**) *In situ* hybridization of *Ahd2* and *Th* transcripts in coronal slices of 6 months old mice from more rostral to caudal, comparing *Dot1l*(+/+), *Dot1l*(*fl*/+) and *Dot1l*(*fl*/*fl*), showing especially in the latter clear reductions. (**E**) Quantitative PCR of dopaminergic and midbrain markers comparing p40 and one year old midbrains assessing mRNA levels of *Th*, *Pitx3*, the rostro/ventral marker *Ahd2*, and the dorso/caudal marker CCK similar as what observed in (**A**) and unchanged between p40 and one year old Dot1l cKO mice. The yellow and orange dots represent *Pitx3*(*Cre*/+); *Dot1l*(+/+) in P40 and 6 months old mice, resp. The light and darker blue dots represent *Pitx3*(*Cre*/+); *Dot1l*(*fl*/*fl*) mice. *Additional statistics*: a one-tailed students t-test was used with * FDR-corrected *p*<0.05.

### Decreased Dot1l leads to reduced synaptic outgrowth with increased mitochondrial respiratory chain transcripts

We have summarized our transcriptomics approaches (**Fig. 4A**). Firstly, we investigated young (E15.75) *Pitx3*(*Cre*/+);*Dot1l*(*fl*/+) and *Dot1l*(*fl*/*fl*), shortly after the loss of the mark by sorting the cells, focussing on heterozygous mice the age of 6 months old. Since the median half-life of transcripts is estimated at 7-9 hours, and those of proteins ~48 hours (36, 37), we reasoned that by FACS sorting neurons ~5/6 hours after global loss of H3K79me2, (no more than ~18 hours after the global onset of global hypomethylation) we would have the best chance finding direct transcriptional roles of Dot1l/H3K79me (**Fig 4B**). Deseq2 analysis showed 535 genes significantly deregulated (**Fig. 4B**). We found only a marginal number of genes overlapping with E14.5 Pitx3 targets (9/269) (**Sup. Fig. 5A/B**). We next performed Panther cellular component analysis (38) and found genes involved in neuronal outgrowth, Wnt-related (Beta Catenin binding), and mitochondrial (mt) genes being deregulated (Deseq2-Adj.*p*-value<0.05 in *Dot1l*(*flox*/*flox*)). Surprisingly, separating total deregulated genes into *up* and *down*-regulated, especially genes related to mitochondrial function were up-regulated, while genes involved in neuronal outgrowth and Wnt-regulation were down-regulated, with the exception of the Wnts, Wnt7a and Wnt7b **(Fig. 4B-E)**. Further analysis of the GO term ‘Synapse’ revealed that the only exceptions were genes involved in vesicle regulation and endocytosis, the large majority of genes related to synapse function were down-regulated (**Fig. 4C**). Analysis of the GO term ‘Mitochondrion’ showed especially genes of the respiratory chain, the mitochondrial inner membrane in general, and the mtRibosome being up-regulated (**Fig. 4D**). Finally, also 27 pseudogenes related to ribosomal, ATPase, and anti-oxidation that related to the encoding group of up-regulated (only 1 down-regulated), with both Sod1 and the Sod1-pseudogene gm8566 as an example (**Fig. 4D/Sup. Fig. 5C resp.**). Taken together, in maturing DA neurons, genes involved in neuronal outgrowth are generally activated by Dot1l and/or H3K79 methylation, while ribosomal and mitochondrial genes of the respiratory chain are repressed (**Fig. 4F**), with Wnt signaling/targets and Wnts as potential intermediators/comodulators (**Fig. 4E, Sup. Fig. 5A/B**).

**Figure 4.**
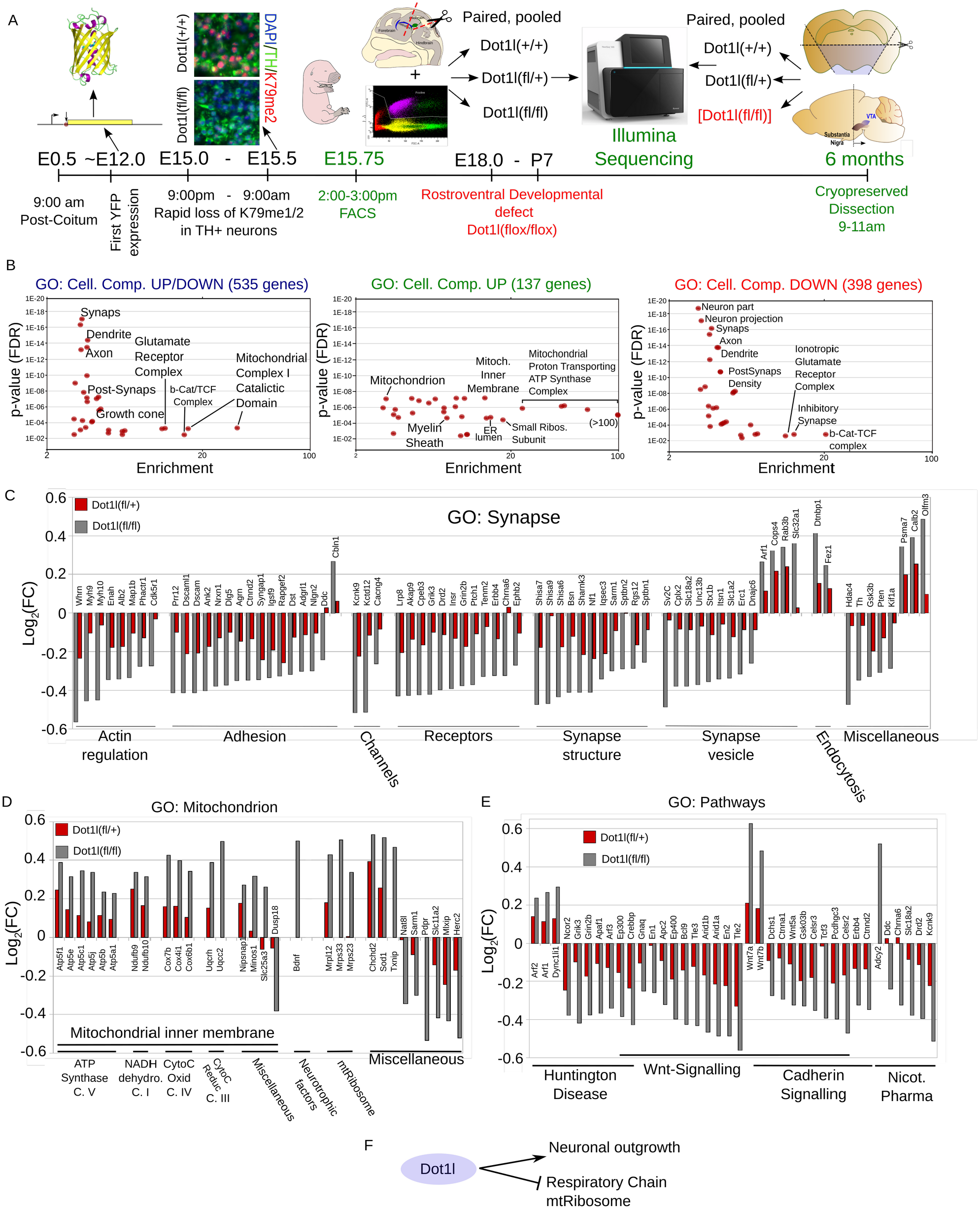
Transcriptome analysis of E15.75 DA neurons with reduced Dot1l/H3K79me2 levels. (**A**) A time-line overview of the set-up. At E12.0 the first YFP expression appears. Between E15.0 (9;00 PM) and E15.5 (9:00 AM) the majority of H3K79me1/2 is strongly reduced. At E15.75 (~3:00 PM) neurons were sorted (FACS) based on Cre induced YFP expression. Per pregnant female, only three embryos (each genotype condition) were used to ensure correct pairing and reduce variation. Per sample, three or four embryos were pooled to generate 3 conditions of n=3 that were used for Illumina sequencing. (**B**) Cellular component analysis (Panther) of in total 535 genes (398 down/137 up) were found deregulated (p-Adj.<0.05) following DEseq2 analysis for differential expression, Dot1l(+/+) vs Dot1l(fl/fl). Total or solely up/down-regulated genes, as a threshold for Panther gene-ontology we have set a minimal enrichment 3 fold and a *p-*value<0.01. (**C**) 2Log(fold change) of differential expressed genes (Dot1l(+/+) vs Dot1l(fl/fl)) of most prominent GO terms found in: Synapse (**C**), Mitochondrion(**D**), and an overview of GO pathways (**E**). In (**F**), an overview of the key postmitotic DA gene-regulatory roles for Dot1l. In DA neurons, at physiological levels, Dot1l activates neuronal outgrowth while repressing ribosomal and mitochondrial transcripts.

### Respiratory chain transcript abundance remains up in 6 months old neurons with reduced Dot1l

Since cKO mice show a cellular developmental defect (**Fig. 3**), we mostly focused on the transcription profile of heterozygous adult *Dot1l* mutants mice. Since technically unbiased DA sorting is limited to prenatal stages, instead, we used a cryopreserved midbrain dissection method with low technical variation and high reproducibility (**Fig. 5A).** With DEseq2 analyses, we found 33 genes and 12 pseudogenes differential expressed (*p*-adj.<0.05) of which only 2 were down-regulated (**Fig. 5B/D**). Panther analysis of these genes showed that especially genes encoding components of the respiratory chain, ribosomal subunits, and mitochondrial regulators were increased (**Fig. 5C**). In line with the E15.75 profile, the 12 pseudogenes (all upregulated) had ribosomal genes, ATPases as parental genes, thus relating to the deregulated set of encoding genes (**Sup. Fig. 6A**). Almost half of the validated genes were mitochondrial, including the down-regulated gene *Guf1* (**Fig. 5B/D**), which is a known regulator of the respiratory chain, though, especially in conditions of cellular stress(39). Next to mitochondrial genes, among which those involved in anti-oxidation, mitophagy, protein syntheses, and mitochondrial carrier, a quarter of the dysregulated genes encoded ribosomal subunits (**Fig. 5D**), suggesting a broad role of DOT1L in mitochondrial and ribosomal gene regulation in adult neurons (**Fig. 5C/D**).

**Figure 5.**
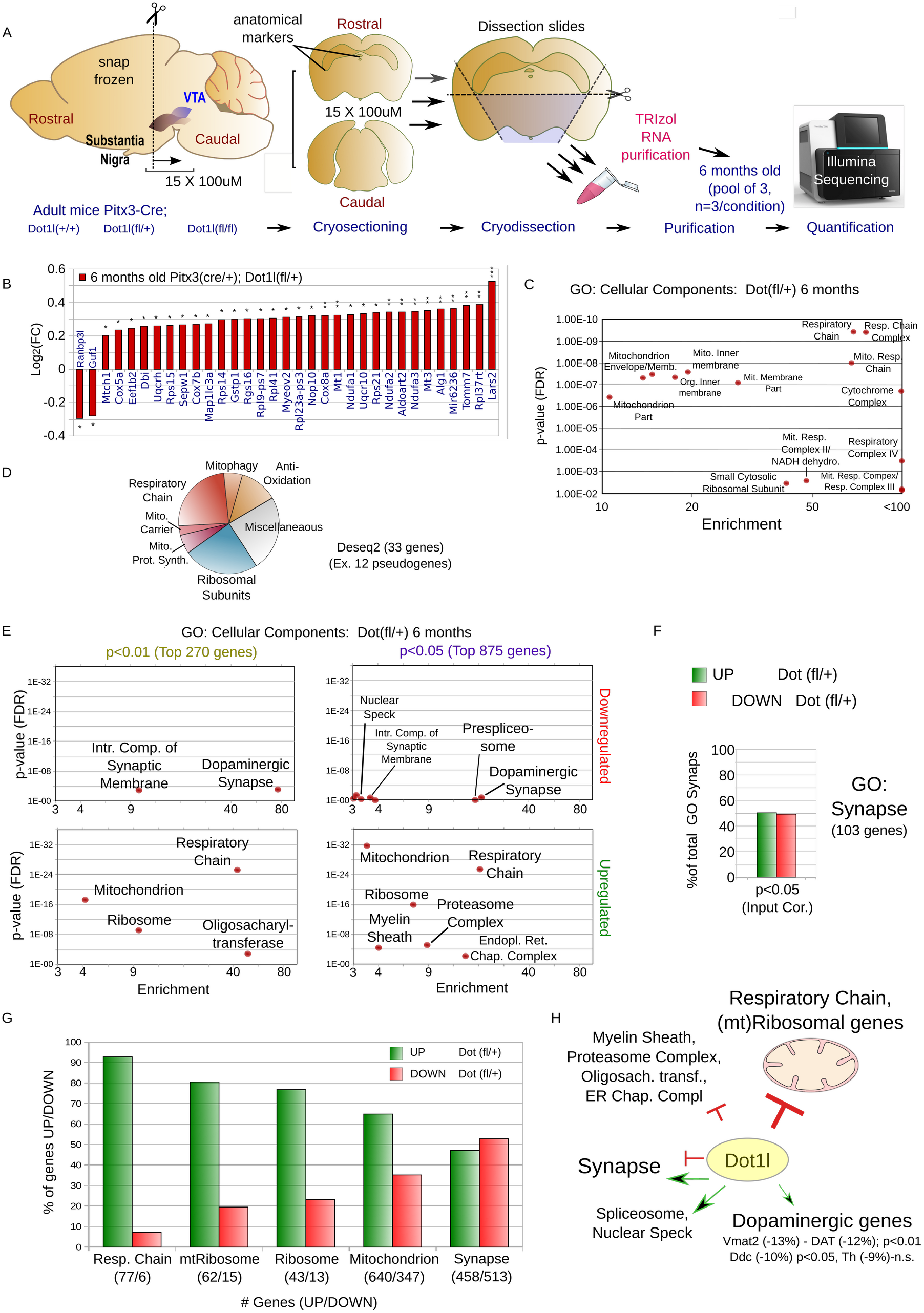
Transcriptome profiling of 6 month old *Pitx3*(*Cre*/+); *Dot1l*(*fl*/+) mice. (**A**) Schematic overview of dissection and transcriptomics approach. Snap frozen 6 months old mice brains were sliced in a cryostat till rostral anatomical markers of the midbrain were reached. Exactly 15 slices of 100uM thick were sliced and the ventral midbrain area was further dissected by removing the dorsal midbrain and hemicortices, collected and thawn immediately into Trizol. Three (n) of paired-pooled samples were generated for each of the 3 conditions (*Pitx3*(*Cre*/+);*Dot1l*(+/+), (+/*fl*) *or* (*fl*/*fl*) and the pooled samples were Illumina sequenced and analyzed following DEseq2 statistics. (**B**) Bar diagram of all DEseq2 p-Adj. P<0.05 significantly deregulated genes presented as Log2 fold change. (**C**) Panther Cellular Component analysis of the Deseq2 differential expressed genes (cut-off: minimal enrichment 3, p<0.01). (**D**) Gene function diagram of all DEseq2 deregulated genes. (**E**) Cellular Component analysis of top *down*-(upper diagrams) and *up*-(bottom diagrams) regulated genes with either a non-adjusted p<0.01 (left diagrams) or p<0.05 (right diagrams). Performed with Panther Cellular Component analysis (minimal enrichment 3, p<0.01). (**F**) Bar diagram representing the percentage of GO: Synapse genes (103 in total) being up or down-regulated when retrieved from p<0.05 (top 875 genes) after correction for the slightly over-represented number of up-regulated genes within this group (376 down/499 up). (**G**) Bar diagram of bottom-up analyses of up versus down-regulation of total RC, mtRibosome, Ribosome, Mitochondrion and Synapse related genes. (**H**) Graphical summary of transcriptomic changes and Dot1l targets in DA neurons with especially respiratory chain genes are up-regulated in heterozygous *Dot1l* floxed DA neurons, suggesting a repressive function of Dot1l. *Statistics*: *** represent DEseq2 Adjusted *p*<0.001, ** *p*<0.01 and \**p*<0.05. In **(E)** GO end leaves were are not shown, e.g. the *respiratory chain* but not the individual complexes.

Since we used dissected midbrains the effect size of deregulated genes ubiquitously transcribed is limited because floxed neurons only comprise a portion of the dissected region. Therefore, we have analyzed also larger groups of 270 (p<0.01, non-adjusted) and 875 (p<0.05, non-adjusted) genes (**Fig. 5E**). These larger groups are withal more equally divided between up/down-regulated genes (**Sup. Fig. 6B**), presumably as a result from especially the upregulated (mitochondrial genes) being the primary group with the largest effect size revealed by Deseq2 adjusted p-values. However, deregulated synaptic genes were not as consistently down-regulated as observed at E15.5 (**Fig. 5F**).

We found very similar outcomes with GO analyses, regardless of the statistical cut-off, mostly upregulated respiratory chain and ribosomal genes (**Fig. 5D-E**). Interestingly, with this GO approach, we did find groups of differential expressed genes involved in myelin sheath formation, the proteasome, and ER chaperones among the upregulated genes (**Fig. 5D-E).** With our approach, the effect size following down-regulation may be slightly underestimated for ubiquitously expressed genes. However, at least this will *not* be so for dopaminergic genes that are both specific to and critical for DA neurons. Especially DA transporters, cholinergic receptors, and genes involved in DA metabolism will have comparable effect sizes between up/down-regulation, though, were only relatively slightly down-regulated, with the rate-limiting DA producing enzyme, *Tyrosine Hydroxylase* (*Th*), not among the 875 significantly deregulated (i.e. the top 376, *p* non-adj < 0.05, down-regulated) genes. In addition, this is in line with our qPCR analysis at P40 and the ISH analyses at P7 and 6 months **(Fig. 3)** and the minor effects observed in the heterozygous *Dot1l*(*fl*/+) at E15.75 (**Fig. 3D**). Opposite to the developmental E15.75 stage, the top deregulated synaptic genes at 6 months heterozygous *Dot1l* DA neurons were slightly more often up-regulated, even after correction for the over-represented number of up-regulated versus-down-regulated genes in this group (**Fig 5F, Sup. Fig. 6B**).

Finally, to further circumvent technical biases, we also analyzed the data set using a bottom-up approach by selecting known genes encoding components of the respiratory complex (RC) and the (mt)ribosome, synapse (GO:0045202) or mitochondrial (MitoCarta 2.0) genes in general. We found the large majority of the RC(93%), mtRibosome(81%) and ribosome(77%) genes up-regulated on average, with even 65% of the total 987 mitochondrion related transcripts retrieved from our data-set (cutoff minimal 80 read-counts) being up-regulated (**Fig 5G**). Of the retrieved synaptic genes (971) was slightly more deregulated towards the downside (53%, **Fig 5G**). In **Figure 5H** the transcriptional roles of Dot1l in DA neurons are summarized.

### mtDNA encoded genes and a selection of genes involved in mitochondrial morphology form exceptions

While RC components, including those coded on the mtDNA (**Fig 6A**) provided that they had passed our experimental pipeline (i.e. ND1,2,4,5;mtCyb;Cox1—*HeavyStrand*, ND6— *LightStrand*), were near completely upregulated at E15.75 (**Sup. Fig. 6C**), as near all mtRibosomal subunits were found to be up-regulated (**Sup. Fig. 6D**). However, in 6 months old mice there are a few exceptions of complex I and complex V members (**Fig. 6A, Sup. Fig. 7A**). In particular, while nuclear-encoded complex I members of the RC and ribosome are still generally upregulated, those encoded at the mtDNA are downregulated in the heterozygous mutant (**Fig. 6B, Sup. Fig. 7B**).

**Figure 6.**
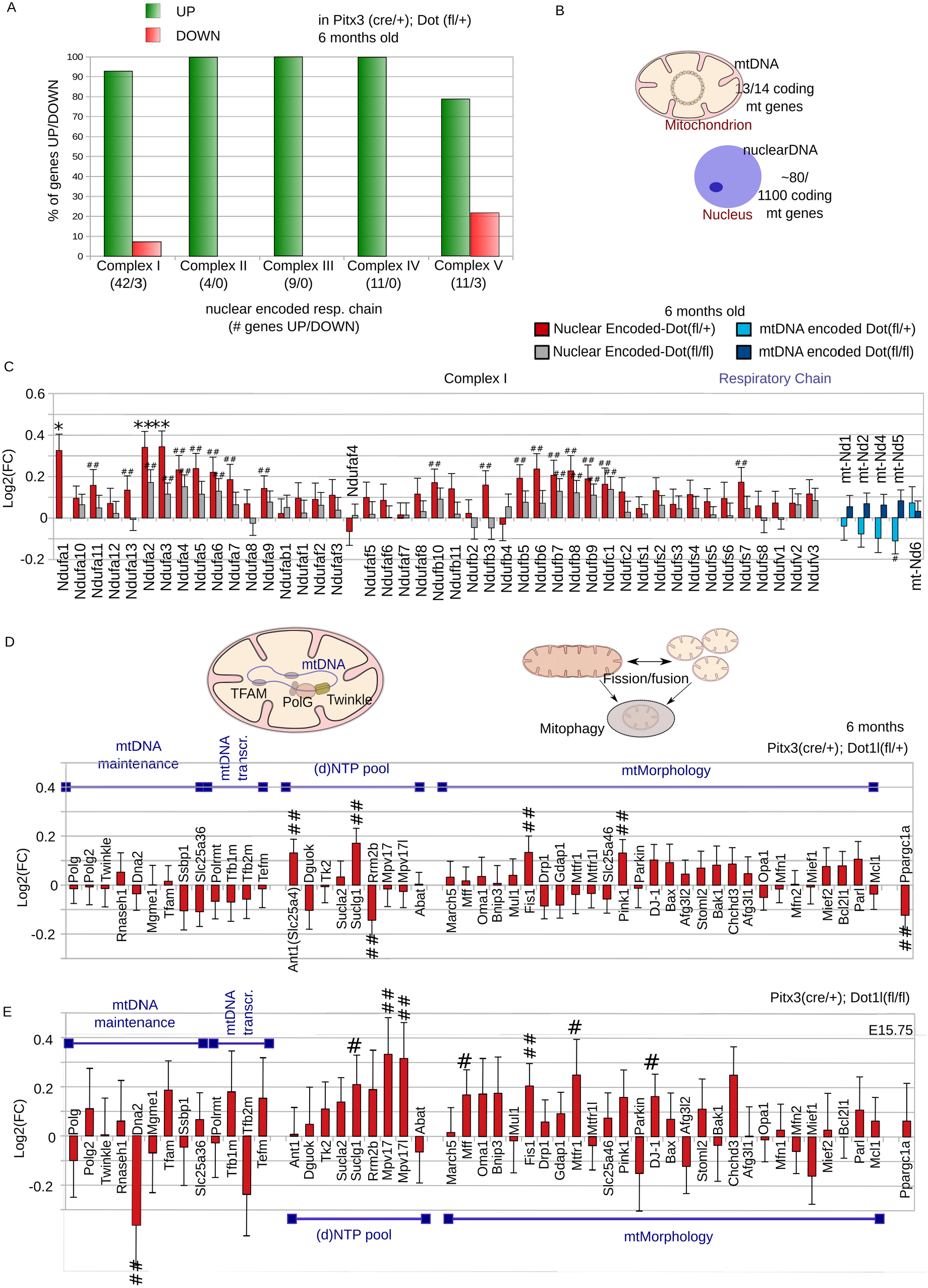
Majority of Respiratory Chain genes, *Pink1*, *Fission*, and *DJ-1* show upward trend following Dot1l reduction. (**A**) Ratio of genes up-versus down-regulated in heterozygous *Dot1l* floxed DA neurons. (**B**) Schematic overview of mitochondrial and nuclear DNA, indicating an approximation of the number of (encoding) mitochondrial genes located at the mtDNA and nuclear DNA. Respiratory Chain genes are encoded both at the mtDNA (13 out 14 encoding genes) as well as the nucleus (~80/1100 mitochondrial genes). (**C**) All retrieved complex I respiratory chain genes of 6 months old mice. For nuclear-encoded genes, red bars represent (*Dot1l*(*fl*/+), gray bars represent *Dot1l*(*fl*/*fl*), and for mtDNA encoded genes: light blue bars represent (*Dot1l*(*fl*/+), dark blue bars represent *Dot1l*(*fl*/*fl*). (**DE**) Selection of genes involved in mtDNA maintenance, mtDNA transcription, maintenance of the (d)NTP pool or mtMorphology, either *Dot1l*(*fl*/+) for 6 months (**D**) or *Dot1l*(*fl*//*fl*) for E15.5 (**E**), red bars represent *Dot1l*(*fl*/+), gray bars represent *Dot1l*(*fl*/*fl*). *Statistics:* ** represent DEseq2, p<0.01 and *p<0.05; ## represent non-adj. P<0.05(Wald-test).

We also included the cKO transcript profile here showing, despite the reduction of neurons, RC genes generally more abundant than controls (**Fig. 6B**).

Prompted by the broad deregulation of mitochondrial genes, we selected a set of known critical regulators related to PD and mitochondrial health, like genes involved in mtDNA maintenance and mtMorphology to seek if a shift in mitochondrial maintenance pathways could be observed (**Fig. 6C/D**). Interestingly, in 6-month-old heterozygous Dot1l cKOs, *Fission1* and *Pink1*, but not Drp1 for instance, all key factors that regulate mitochondrial morphology, mitophagy, and import of transcripts into mitochondria (40), were up-regulated (p0.01, non-adjusted) (**Fig. 6D**). Instead, *Pgc1-α* (*Ppargc1a*), a key regulator of mitochondrial respiratory genes and activator of mitochondrial biogenesis was among the top 270 deregulated genes (p0.01, non-adjusted) (**Fig. 6C/D**) (41). *Pgc1-α* seems not down-regulated at E15.75 in cKOs (**Figure 6E**), potentially being down-regulated as a consequence of mitochondrial deregulation in an attempt to compensate for the reduction of DOT1l.

## Discussion

Dynamic epigenetics and transcript profiles may be pivotal in age-related diseases that involve the loss of neurons and their functions. Here, we have studied DOT1l and H3K79 methylation in human and mouse midbrains, and specifically dopaminergic neurons, in an attempt to study their potential as epigenetic and gene-regulatory key players in neuronal aging and disease processes. DOT1l has been indirectly coupled to aging-related processes by its association to epigenetic mechanisms that involve Sirt proteins, known regulators of bioenergetics (15, 42–44). More recently, data suggests that Dot1l itself may be prolonged deregulated in neurons following early life stress (23). Our findings further support the idea that 1) H3K79 methylation can be highly dynamic in neurons, especially in maturing neurons, 2) that transcription of Dot1l is regulated by energetic and oxidative challenges in neurons, 3) that Dot1l is a master regulator of mitochondrial and ribosomal genes in dopaminergic neurons at physiological levels, and, finally, 4) that in the aged human midbrain and substantia nigra (SNc) accumulation of lipofuscin (LF) consistently correlates with H3K79 hypermethylation.

### The relation between Dot1l and H3K79me2 deregulation and cellular stress

Although evidence for a causal role in the deterioration of neuronal function or PD is scarce, LF has been related to lysosomal dysfunction, ER stress, mitochondrial turnover and oxidation, and is generally accepted as an aging hallmark that may play a role in early stages of Lewy body formation (29, 45). Our findings combined, we hypothesize that increased levels of oxidation may increase Dot1l in neurons. However, this may not lead to increased total levels of H3K79me2/3 immediately, as counteracting mechanisms could accelerate too, either as a consequence of increased histone turnover or via the upregulation of demethylating enzymes.

In neurons accumulating LF, a constant state of oxidation maybe registered and signaled by cell-intrinsic pathways, causing a shift towards a (more) catalytic state that also links with mitochondrial dynamics. We can see Dot1l as an intermediate of this process. It seems likely that the observed H3K79 hypermethylation represents a chronically shifted epigenetic equilibrium, potentially being more harmful instead of guarding cellular homeostasis. Activation of mechanisms that are normally effective to counteract pathogens, as has been shown by roles for Dot1l in viral responses, may damage the cell following long-term overactivation.

Neurons that do not succeed breaking down aggregates and (macro)vacuolar structures may chronically register a state of oxidation. We can imagine that high levels of H3K79me2 that associate with LF are the end state of such a process of, among other effectors, chronically increased Dot1l levels.

Reduced counteracting mechanisms, for example, a reduction of histone turnover following reduced H3.3 expression could further add to the shift in methylation states, with H3K79me2 residing on the portion of H3.3 subunits that may turnover most rapidly in younger neurons, a process that has been suggested to influence the H3K79 methylation state as well (2, 46). Moreover, histone turnover can be a more prominent effector at histones marked by active marks, which could explain why H3K27me3 does not accumulate in LF-associated nuclei. However, if we compare the rapid loss of post-mitotic neuronal H3K79me2 following floxed *Dot1l* alleles, we would not be surprised if demethylation enzymes, like the recently described H3K79 demethylase KDM2B (47), further add significantly to the equation. At least, the fast global turn-over of H3K79 methylation in post-mitotic neurons suggest different dynamics than H3K27me3 which takes approximately 3-4 weeks after cKO of methyltransferase activity (31).

Another point worth to discuss is the divergent and, on average, only mild increase of H2K79me2 in Parkinson’s Disease (PD), often lower than surrounding LF containing neurons or compared to TH neurons that contain LF. Whether the two individuals with higher levels in TH+ cells are exceptions with an atypical form of PD, simply suggesting that H3K79 hypermethylation is a rare phenomenon in DA neurons, or if most SNc DA neurons are too vulnerable to survive (or express TH) with a substantial LF/H3K79me2/3 burden, remains uncertain. Interestingly, in aging rhesus monkeys, LF associates especially with VTA DA neurons and less with those of the SNc (33). This could be the consequence of subtypes of DA neurons, mostly found in the VTA, still expressing TH while accumulating LF. Since we have found many LF encircled large nuclei in the middle of the substantia nigra, surrounded by TH+ neurons with similar large nuclei and macro morphology with clusters and similar relative distance (observational), we can imagine that (some) of the strongly LF+ neurons used to express TH. It would be interesting to study the relationship between TH levels and LF or lysosomal defects in more depth since in an ultimate attempt to clear neurons from dysfunctional organelles and aggregates may undesirably affect functional proteins like TH. Another explanation would be that SNc DA neurons develop LBs instead of LF in times of stress, perhaps as a consequence of the large number of mitochondria, of which (near-)complete units form main components of matured LBs (29).

### The potential of Dot1l as a therapeutic target to regulate neuronal mitochondria

The broad down-regulation of nuclear encoded respiratory genes is a transcriptional hallmark of PD (48). However, attempts to improve (mitochondrial) homeostasis in PD-models by over-expression of PGC1-α (Ppargc1a), a key inducer of mitochondrial biomass, have not been successful and may even worsen the pathological state (49). However, in several ways manipulation of mitochondrial transcription through Dot1l could differ from PCG1-α. Firstly, whereas hypothetically PGC1-α needs to be overexpressed, Dot1l would need to be downregulated. The former has been suggested to result in several off-target effects at non-physiologically high levels(49). Secondly, depletion(cKO), but not reduction(heterozygous), of Dot1l resulted in a defect but did not progressively seem to harm the survived population. Third, sub-physiological levels of Dot1l relate to slightly lower PGC1-α, perhaps increasing respiratory chain and anti-oxidation transcription and innermembrane-complex-turnover in the absence of increased mitochondrial mass or fusion. Moreover, genes related to the proteasome were increased, which may alleviate the burdens of cellular waste. If neurons increase proteasomal protein complex specific degradation and turnover of mitochondrial complexes, the chance of aggregate formation could be reduced reducing last-resort pathways, ending up as disease-associated macro-vacuoles with LF and LBs. In summary, targeting DOT1L has the potential of being a more subtle and specific target than PGC1-α worth to be further investigated.

Finally, another interesting broad regulator of RC genes is the mitochondrial translation elongation factor GUF1, the only mitochondrial factor that is significantly reduced at 6 months (**Fig 4B**). Ablation of the highly conserved *C. elegans* homolog of GUF1, mtEF4, protects against paraquat toxicity in *C. elegans* (50). Next to this, the conserved yeast GUF1 regulates the RC and ribosomes under stressed conditions (39).

### Transcriptional exceptions: mtDNA encoded respiratory genes

Although few transcriptional or epigenetic mechanisms seem able to compensate completely for the transcriptional consequences of Dot1l reduction, the two paralogs Atp5g1 and Atp5g3 form exceptions involved in complex 5 abundance (**Sup. Fig. 7A,B**). This is also the case for mtDNA encoded genes that seem to be decreased in the heterozygous cKO while even slightly increased on average in the *Pitx3-Cre*; *Dot1l*(*fl*/*fl*) full cKO (**Sup. Fig. 7B**). Remarkably, a similar discrepancy between nDNA and mtDNA transcriptional deregulation has been observed in a study performed by NASA on the health of astronauts during spaceflights that was recently published (51). Cells that underwent a period of in space showed nDNA respiratory transcripts were broadly reduced, similar to PD (41), while mtDNA encoded transcripts were upregulated (51).

### Regulation of mitochondrial turnover and mitogenesis

If no transcriptional compensation of mitochondrial regulation abnormalities can be found then increasing the turnover of mitochondrial components, or whole mitochondria may be another way for cells to avoid a mitochondrial overload following Dot1l dependent ‘mtComponent’ deregulation. Indeed we find regulators of mitochondrial morphology and turnover, like *Fis1*, *Pink1*, and *Dj1* (40), up-regulated both at E15.75 and 6 months (non adj.p<0.5/p<0.1) (**Fig. 6D-E**), as well as proteosomal components. This contrary to the stress-regulated mitophagy pathways that include *Drp1* (Ganesan et al., 2019), which is slightly up-regulated on average at E15.75, but down-regulated at 6 months.

In summary, Dot1l and H3K79 hypermethylation are regulated by and associate with states of neuronal stress in human neurons. The fast turnover of H3K79 methylation and dynamic regulation of Dot1l transcription in neurons suggests an adaptable mechanism with Dot1l as central master repressor of mitochondrial transcripts. In addition, DA neurons can survive following complete Dot1l/H3K79me depletion, without any sign of a progressive phenotype, putting Dot1l forward as a potentially better therapeutic candidate then PGC1-a in order to enhance transcript abundance of OXPHOS genes. For this purpose, we suggest that the roles of the various methylation states and methylation independent neuronal functions of DOT1l and H3K79me1/2/3 in neurons will need to be dissected in more detail.

## Methods

### Human tissue and anatomical selection

Human donor specifications are provided in **Supplemental Table 1** (‘Experiment 1’: **Fig. 1A-D, G**) and Supplemental **Table 2** (‘Experiment 2’: **Fig. 1 E,F**) and were obtained from the Netherlands Brain Bank (NBB) as paraffin blocks of hemi-midbrains (**Sup. Fig. 3A**). The rostral to caudal tissue sections were collected from two or three adjacent paraffin tissue blocks for each donor (as provided by the brain bank). Slices were matched by determining the position where the first most rostral, low vulnerable TH+ VTA neurons flanking the dorsal side of the red nucleus using TH-DAB stainings (**Sup. Fig. 3B/C**). For experiment sections proximate, but caudal to this position, have been used retrieved from one mid-DA system paraffin block for each donor.

### Immunohistochemistry human brain tissue

Deparaffination/rehydration was performed in: 3×4’ in xylene, 2×4’ 100% ethanol, 2×4’ in 96% ethanol, 4’ in 80% ethanol, 4’ in 70% ethanol and 4’ in 50% ethanol. Then slides were washed twice for 5 minutes in PBS and incubated for 5 min in 0.01M citrate buffer at RT. Antigen retrieval was performed in 0.01M citrate buffer by heating samples for 2 min at 800W approaching boiling point. Subsequently, an 85-93 degrees Celsius temperature was maintained by giving short 200W bursts for 20’ with increasing interval. Slides were subsequently incubated in a 67 degrees stove for 60’. After cooling down the slides in a water bath they were washed in TBS for 5’ and then blocked for 60’ with 5% NDS in THZT. Slides were washed 2×5’ in TBS and then Incubated in sheep-α-TH (1:500, Milipore, Ab1542) overnight. Then slides were washed 3×5’ in TBS and incubated in donkey-α-sheep-488 (1:500, Life Technologies, A11015) in TBS for 90’. Slides were subsequently washed 3×5’ in PBS and blocked in 5% FCS in THZT for 60’. Slides were then washed again in 2×5’ PBS and incubated overnight in rabbit-α-H3K79me2/3 or H3K27me3 in THZT. Subsequently, slides were washed in 3×5’ PBS and incubated in goat-α-rabbit-555 (1:1000, Life Technologies, A21207) in PBS for 90’. Slides were then washed in 2×10’ in PBS and subsequently incubated in DAPI (1:3000) in PBS for, 5’ in experiments 1, 8’ in experiment 2 (optimized for large neuronal nuclei). Finally, one PBS wash for 5’ and embedded in Fluorsave (Millipore, 2848323).

### Imaging

Microscope imaging for analysis of IMHC experiments was performed using a Leica DFC310 FX microscope with Leica MM AF 1.6.0 software. For human tissue, 2×2 or 4×4 images were generated at a 40x magnification of the SNc for each slice. For experiment 1, to asses rostral to caudal LF associated nuclear staining, 10 images were generated from three separate tissue blocks originating from a 79 year old donor. For the additional donors 2 Imagesoriginating from 2 slices from the 90 year old and 77 year old donor. In experiment 2, images generally contained 5-50 TH+ neurons per image from lateral to medial. 2×3 images were generated from two slices separated by 160um and spread from lateral to medial. VTA DA neurons were captured in small groups or single neurons surrounding the red nucleus since they are more scattered over a larger area. The location and subdivision of the VTA area was based on Root and colleagues (52).

### Quantification of IMHC signal humane tissue

Quantification of H3K79me2/3 staining in Th+ DA neurons was performed using ImageJ 1.51j8 and blinded (imaging and quantification were independently and blinded performed by two individuals). Reference nuclei and the nuclear area was determined by converting the DAPI signal to binary images, and selecting nuclei regions of interest (ROIs) at size (2000-6000 pixels) and circularity>0.7, selecting and labeling the majority nuclei in experiment 2. Next, the regions of interest were overlaid with color combined to separate DA (TH+) with/without inclusions or LF associated neurons. For experiment 1, LF+ nuclei ROIs were manually selected and, opposite, manually deselected from the reference nuclei set. ROIs were overlaid with 16-bit H3K79me2 image layers to quantify the average signal in each nucleus. In experiment 2 this was followed by a color combine image overlay to select DA neuronal nuclei neurons, those with/without LF or black pigmented inclusions. H3K79me2 levels of LF+ were manually validated in experiment 2. We found levels of H3K79me2 low nuclei were lower than background of some fiber structures that were withal not equally divided between and within pictures, which prompted our decision to perform calculation in represented in figure 1 without background correction. In **Supplemental Figure 2**, the individual measurements with/without local background correction have been presented as well (which had no statistical consequences). For this, we have selected >10 area’s of each photo, in between cells in low nuclear regions but avoiding fiber tracts depleted from cells but with high local levels of background to determine the average background for each photo which were used for individual background correction (IBC) by subtraction. As a comparison, an equal background correction (EBC).

### Animals

The C57BL/6J mouse line was used as background strain (Charles River). Mice expressing Cre from the *Pitx3* locus and Cre induced YFP from the *Rosa26* were previously described (35) and crossed with floxed *Dot1l* mice that were a kind gift of Dr. Zhang (34). *Pitx3*(+/+); *Dot1l*(*fl*/+); *Rosa26*(*YFP*/*YFP*) breeding does and *Pitx3*(*Cre*/*Cre*); *Dot1l*(*fl*/+); *Rosa26*(+/+) breeding studs were crossed to generate offspring nests containing all experimental genotypes. All procedures were according to and fully approved by the Dutch Ethical Committees for Animal Experimentation (University of Amsterdam).

### Immunohistochemistry mouse brain tissue

PFA sucrose preserved brains were fixed as whole heads for 4 hours (E16 or younger), otherwise as whole brains for 7 hours, in 4% paraformaldehyde (PFA) at 4C, put in PBS containing 30% sucrose(w/v) until they sunk, and kept at −80 till sliced at 16μm sections using a cryostat. To perform IHC the slides were thawed, washed with PBS(3X), post-fixated with 4% PFA for 10 minutes, and washed with PBS(3X) again. Using antibodies against histone marks, antigen retrieval was performed by incubation for 5 min in 0.01M citrate buffer at RT, heating in 0.01M citrate buffer by heating samples for 2 min at 800W in the microwave to +/− 90 degrees, followed by 200W pulses for 20’ and incubated at 67 degrees for 1 hour. After cooling the slides down they were blocked with 5% NDS in THZT, washed in TBS(3X), and incubated in sheep-α-Th overnight. The next day, they were washed with TBS(3X) before incubation with donkey-α-sheep-488 in TBS for 90 minutes, washed in PBS(1X), blocked in 5% FCS in THZT for 1 hour, washed in PBS(2X) and incubated o/n with rabbit-α-H3K79me1/2 in THZT. The following day, slides were washed in 3×5’ PBS and incubated with Goat-α-rabbit-555 in PBS for 90’. Slides were then washed in 2×10’ in PBS and subsequently incubated in DAPI in PBS for 7’. Finally, slides were washed once again in PBS for 5’ and embedded with Fluorsave (Millipore, 2848323). Microscope imaging for analysis of IMHC experiments was performed using a Leica DFC310 FX microscope with Leica MM AF 1.6.0 software.

### Primary neuronal culturing and treatment

12 wells plates were coated o/n with poly-L-lysine (0.1 mg/ml [SigmaAldrich]) in 0.1 M borate buffer pH 8.5 (0.04 M Boric acid [Merck Millipore], 0.01 M Borax [SigmaAldrich]). 1 ml of plating medium (MEM Eagle’s with Earle’s BSS medium [Invitrogen], supplemented with 10% heat-inactivated FBS [Gibco], 0.45% (wt/vol) glucose [Merck Millipore], 100 mM sodium pyruvate [Invitrogen], 200 mM glutamine [Invitrogen] and penicillin/streptomycin [Invitrogen]) was added to each well and the plates were stored in the incubator on 37°C with 5% CO2. Primary cortical cultures were prepared from E17.5 C57BL/6 mice. E17.5 cortical tissue was dissected in ice-cold dissection medium (HBSS medium [Invitrogen], supplemented with 100 mM sodium pyruvate [Invitrogen], 20% (wt/vol) glucose [Merck Millipore] and 1 M Hepes [SigmaAldrich]) washed and trypsinized at 37°C for 20 min and 100 μl of DNase [Thermo Fisher Scientific] was added for 5 minutes before dissociation by trituration. 1Ml plating medium containing 100.000 cells/ml were added to each well of a 12-wells plate containing 1ml of pre-incubated medium. After 2 days, 50% of the medium was replaced with maintenance medium (Neurobasal medium [Invitrogen] supplemented with 2% B27 [Invitrogen], 200 mM glutamine [Invitrogen] and penicillin/streptomycin [Invitrogen]) and 10 μM 5-Fluoro-2’-Deoxyuridine (Fudr, [Sigma-Aldrich]). At DIV5 and DIV11 50% (1ML) of the maintenance medium was replaced containing a vector, 200 μM of AICAR or 20μM of H2O2, to reach a final concentration of resp. 100 μM and 10μM.

### In situ hybridization

Mouse brains were immediately snap-frozen using dry ice. *In situ* hybridization was performed as previously described (Ahd2, (53); Th, (54))

### Antibody list

| Antibody | Product number | Concentration |
| --- | --- | --- |
| rabbit-α-H3K79me2 | Abcam, Ab3594 | 1:1000 in THZT |
| rabbit-α-H3K79me3 | Abcam, Ab2621 | 1:750 in THZT |
| Rabbit-α-H3K27me3 | Millipore, 17-622 | 1:1000 in THZT |
| sheep-α-TH | Millipore, Ab1542 | 1:500 in THZT |
| Goat-α-rabbit-555 | Life Technologies, A27039 | 1:1000 in PBS |
| donkey-α-sheep-488 | Life Technologies, A11015 | 1:500 in TBS |
| chicken-α-GFP | chicken-anti-GFP | 1:1000 in THZT |
| rabbit-α-H3K79me1 | Abcam, ab2886 | 1:100 in THZT |
| Goat-α-chicken-488 | Thermo Fisher, A-11039 | 1:1000 in PBS |
| Donkey-α-rabbit-594 | Thermo Fisher, A-21207 | 1:1000 in PBS |

### Fluorescence Associated Cell Sorting (FACS)

Dissected midbrains were dissociated using a Papain (Worthington) and neurons were sorted using a BD FACS Aria III cell sorter. Gates were set on forward scatter versus side scatter (life cell gate), on forward scatter versus pulse width (elimination of clumps), and on forward scatter versus fluorescence channel 1 (528/38 filter; GFP fluorescence: sufficient to sort YFP fluorescent neurons). Cells were sorted with an average speed of ~5000 cells/second into a drops of sterile PBS and collected in Trizol LS reagent (Invitrogen) ratio 1:3 in a precooled collection chamber at 4C, collected, mixed well and stored till further use at −80C.

### RNA Sample generation

RNA from FACS isolated cells was purified using a Trizol LS (Sorted neurons, Invitrogen) according to the manufacturers protocol (3 volumes of PBS/cells:1 volume Trizol LS). For adult animals, brains were snap-frozen on dry ice, sliced in a cryostate till the anatomical markers that indicate the rostro-diencephalic DA area. Subsequently, 15 slices of 100um were sliced, the DA area was dissected, thawed and lysed directly in 0.5Ml TriZOL (Invitrogen) and stored at −80C till further use. Three samples were pooled (using only one paired set of +/+, fl/+, and fl/fl from each nest/uterus) that were subsequently cleaned and DNAse treated using the zymo-clean&concentrator kit (Zymo Research). For primary neurons, 0.5 ml per 12 wells well was used.

### RNA sequencing

The quality of the RNA samples was determined using a Fragment Analyzer, samples with an RQN score > 6 passed QC. The NEBNext Ultra Directional RNA Library Prep Kit for Illumina was used to process the samples according to the protocol NEB #E7420S/L (New England Biolabs) to fragments of 300-500 bps. Clustering and DNA sequencing was performed using the Illumina NextSeq 500 device according to manufacturer’s protocols and with a minimum depth of 18 million quality-filtered reads per sample. The read counts were mapped against Mus musculus GRCm38.p4 and paired analyzed using DESeq2 to determine differential expression between groups (Genomescan BV).

### Quantitative PCR

Total RNA was purified by applying Trizol (Invitrogen) to cells according to the manufacturer’s instructions. qPCR amplification was performed on a *Roche* light cycler using OneStep qPCR SYBR green kits (Qiagen) according to the manufacturer’s protocol. 10 ng total RNA was used as input.

List of primers used:

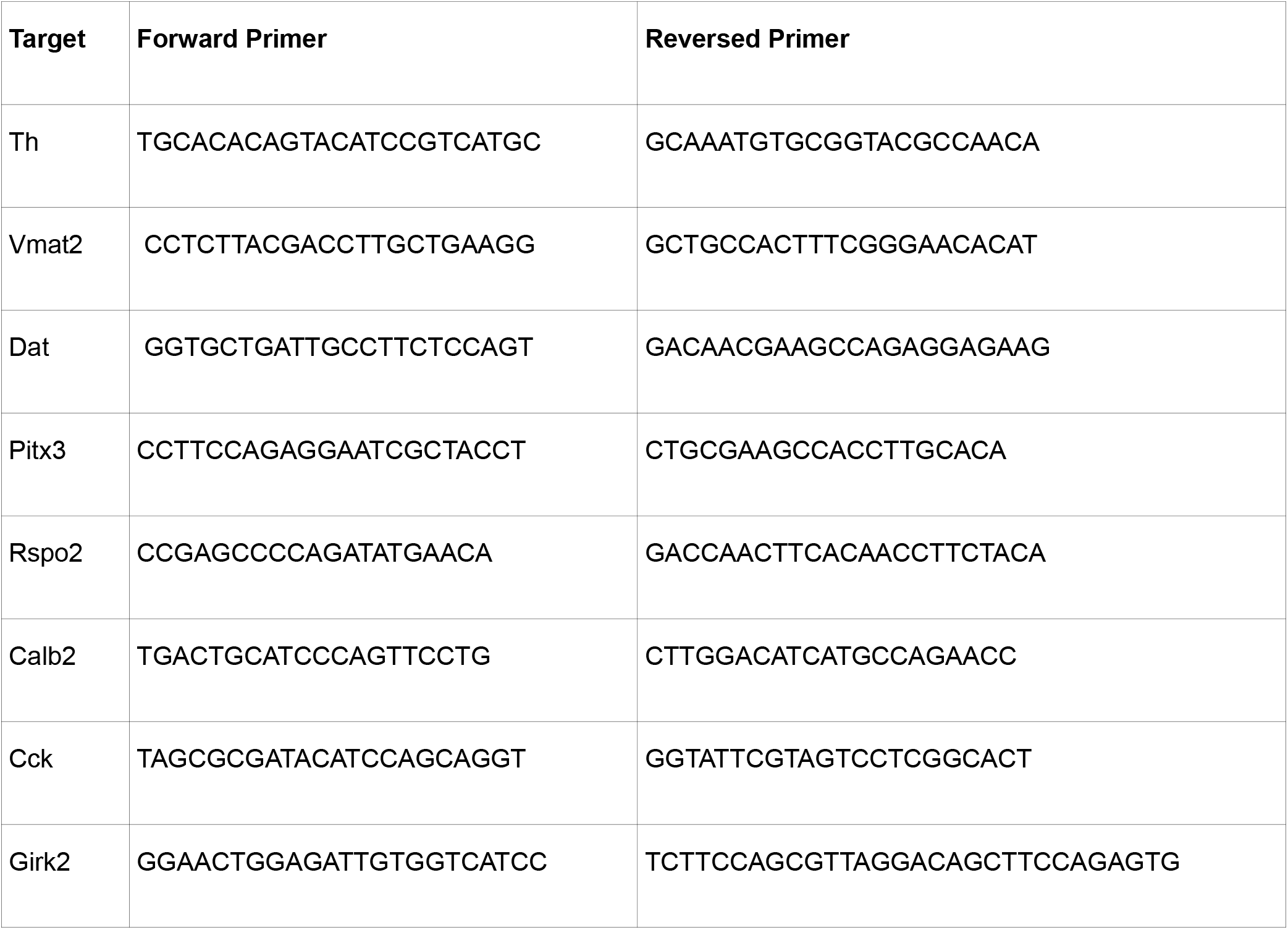
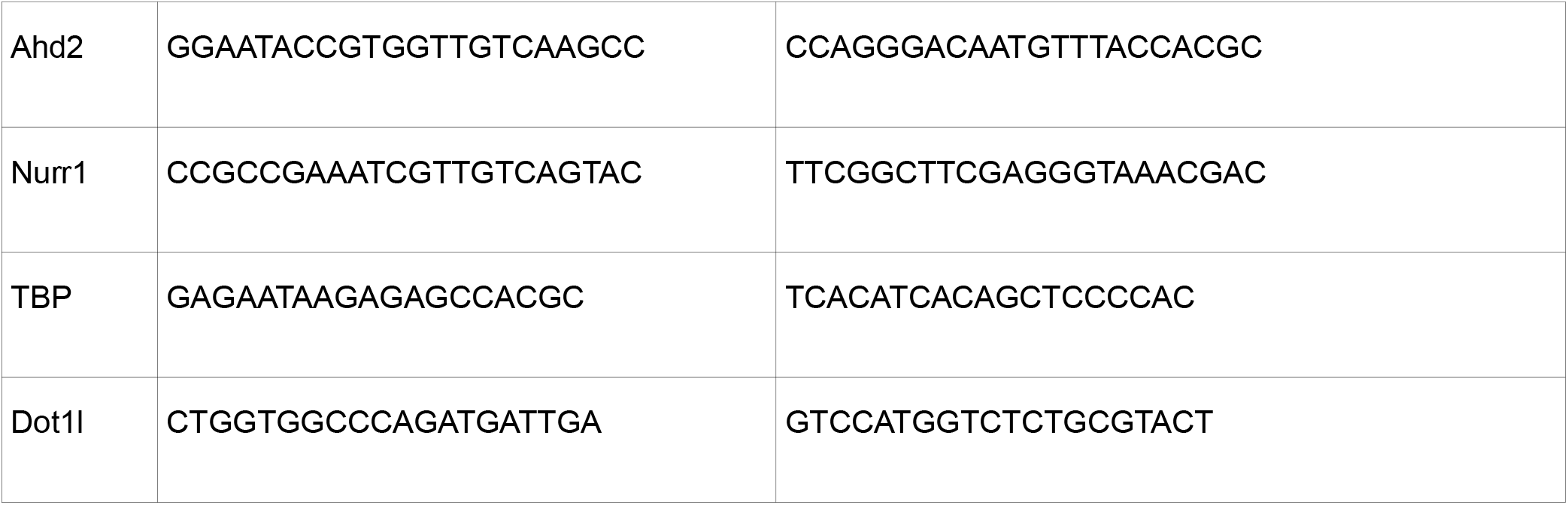

### Statistics and graphs

For one way analyses of the distribution of multiple (4) groups of nuclear H3K79 staining intensity in Figure 1, we have used a Kruskal-Wallis non-parametric test https://www.socscistatistics.com/tests/kruskal/default.aspx. To test if the distribution between two non-parametric groups differed Figure 1E (VTA/SNc comparison) we used Mann-Whitney analyses: https://www.socscistatistics.com/tests/mannwhitney/default2.aspx, followed by Bonferroni correction for multiple testing. For data plotting, we have used in figures 1, 3, and 4 https://huygens.science.uva.nl/PlotsOfData/ by Dr. Joachim Goedhart. To analyse the RNA sequencing data for differential expression DESeq2 statistical analyses was used. For other statistics Student’s T-Ttests with parameters as stated in the legends were used.

### Study approval

All animal procedures were according to and fully approved by the Dutch Ethical Committees for Animal Experimentation (University of Amsterdam). All human studies were fully approved by the ethical guidelines of the Netherlands Brain Bank.

## Author contributions

Study and experimental design and writing was performed by H.v.H and M.P.S. IHC and Microscopy of the human studies were performed by H.v.H, L.v.O and P.d.V., quantification was performed by P.d.V and H.v.H. qPCR of neuronal cultures was performed by C.W. The primary cultures were generated by H.v.H. with help of C.W., and L.v.O. All other experiments, analyses and figures were generated by H.v.H. The article was written by H.v.H and M.P.S.

## Acknowledgments

This work was sponsored by the NWO-ALW (Nederlandse Organisatie voor Wetenschappelijk Onderzoek-Aard en Levenswetenschappen) VICI grant (865.09.002) awarded to MPS. The funders had no role in study design, data collection and analysis, decision to publish, or preparation of the manuscript. We would like to thank the Netherlands Brain Bank and Prof. dr. Paul Lucassen for providing human brain samples. We would like to thank Dr. Joachim Goedhart and r. Marten Postma for generating and maintaining the PlotsOfData tool. We would like to thank Dr. Whenzeng Zang for providing us with the Dot1l-loxp mice and Andrea Heredero Berzal for assistance with culturing primary neuronal cultures. Transcriptome data is submitted to GEO: GSE184901 and GSE195257.

## Supplemental Legends

**Supplemental Figure 1 Correlation between lipofuscin and H3K79 hypermethylation.** Rostral to caudal fluorescent microscopy images additional to Fig. 1, of LF autofluorescence associated to large nuclei in the human midbrain of a 79 y/o male donor showing ER lysosomal/lipofuscin structures surrounding the nuclei relating to H3K27me3 (A) or H3K79me3 (B) staining with white arrows indicating large nuclei with low to almost absent H3K27me3 staining. (C) Fluorescent microscopy images of LF autofluorescence associated to large nuclei in the human midbrain of a 77 and 90 y/o. For the latter the white box depicted a few exceptions of relatively low H3K79me2 nuclear levels in the mids of LF associated nuclei with H3K79me2 hypermethylation. In (D), quantified levels of nuclear H3K79me2 are presented, comparing between nuclei that are surrounded by LF (yellow dots) and surrounding reference nuclei (blue dots). (E) Th (green), DAPI and H3K79me3 staining in 90-year-old asymptomatic control shows low to moderate H3K79me3 in aged Th+ SNc DA neurons. Statistics: Mann-Whitney test was performed for each separate section to test if H3k79me2 was increased in lipofuscine compared to reference nuclei. **** represents a p-value<0.0001.

**Supplemental Figure 2 Overview of single nucleus H3K79me2 levels between individuals and cell types.**

For each donor (A-O), we have represented an image of TH+ neurons representing average levels of H3K79me2 staining intensity (a.u.) (upper panel) and each measure point (lower panel). The title colors and ages correspond to different Braak stages (which did not relate to K79me) levels. To visualize the independence of relative intra sample differences to local background, we have shown samples corrected for the local background, while the left panel are uncorrected data corrected for an individual background correction (IBC, described in ‘Methods-quantification’) on the left and corrected with an equal (no) background correction for all samples (EBC) on the right. From left to right, levels corresponding to non-DA nuclei (TH-negative (TN), blue), TH+ nuclei (TH+, dark green), and TH+ nuclei split into nuclei without clear pigmented inclusions (A-, light green) or with clear pigmented inclusions (red, +A). Lipofuscin positive neurons (LFn), were added as 5th condition for each subgraph (Yellow, L). Statistics: Performed on IBC measurements. Mann-Whitney analyses comparing between TH- and TH+ nuclei and between pigmented inclusions and lowly pigmented TH+ neurons adjusted with a post-hoc Bonferroni correction for multiple testing. **** represent p<0.0001, *** p<0.001, ** p<0.01, *p<0.05 and n.s.: non-significant.

**Supplemental Figure 3 human midbrain material overview**

(A) Example of paraffin donor material supplied by the Netherlands Brain Bank with (B) a coronal section stained with anti-TH DAB of a 90 year old donor. (C) Magnifications of TH+ cells in the dorsal VTA, the medial and lateral SNc (C).

**Supplemental Figure 4 Loss of H3K79 mono/dimethylation in post-mitotic dopamine neurons after ablation of Dot1l.**

(A) Sections increasingly lateral from the retroflex and (B) the most medial section of the E15.5 midbrain were studied to further investigate the loss of H3K79me2 following Dot1l depletion in dopamine neurons. In green Tyrosine Hydroxylase positive (TH+) cells were marked together with staining nuclei H3K79me2 (red). Blue arrows indicate several Th+ neurons that still possess clear H3K79me2 nuclear staining, especially increasing when getting further lateral from the retroflex (A) and in the most medial sections of the midbrain (B) Blue arrows indicate two slightly H3K79me2 positive nuclei of TH+ cells. In (C) is a schematic overview of a E15.5 coronal midbrain section represented. The red dotted line represents the cutting position. The red box represents the region shown in (D). In (D) the loss of H3K79me1 is shown, supporting a global loss of H3K79 methylation in cKO Dot1l mice (lower panel), compared to controls (upper panel; Pitx3(Cre/+)). TH is shown in green, DAPI in blue and H3K79me1 in red. The white arrows indicate internal controls of non TH+ cells still clearly positive for H3K79me2. (E) Coronal sections of the left ventral midbrain of E18.5 Pitx3(Cre/+);Dot1l(+/+); Rosa26(YFP/+) (Left-half) and Pitx3(Cre/+);Dot1l(fl/fl);Rosa26(YFP/+) (right half) showing the absence of H3K79me1 (red) in nuclei (blue) of DA neurons (green; TH+) several days after the primary loss has been observed, suggesting Dot1l to be the sole factor to maintain global H3K79 methylation. White arrows indicate TH-negative cells within the SNc that possess normal H3K79me2 nuclear staining as internal controls (Floxed cells in this region are TH+ without exceptions).

**Supplemental Figure 5 Dot1l deregulated genes relate to deregulated pseudogenes but do not overlap with E14.5 Pitx3 (GFP/-) deregulated genes**

(A) Overlay between differential expressed genes comparing E14.5 sorted neurons of Pitx3(GFP/+) versus Pitx3(GFP/-) (Unpublished data) and Pitx3(Cre/+);Dot1l(+/+) and Dot1l(fl/fl). Out of the Pitx3(GFP/-) deregulated genes (263 in total) only 22 overlapped, of which only 6 up- and 3 down-regulated genes were differential regulated in equal directions (up/down). (B) Ranking of co-deregulated genes based on their p-value rank in either the Pitx3(GFP/+) (first column) or Dot1l(fl/fl) (second column) or their rank in the Dot1l(fl/+) (‘level-dependent’) comparison (third column). (C) List of functions related to pseudogenes parent genes, of pseudogenes, found deregulated with Deseq2 analysis.

**Supplemental Figure 6 Dot1l top deregulated genes are upregulated at 6 months and overlap with deregulated pseudogenes**

(A) List of functions related to pseudogene parent genes to pseudogenes found deregulated with Deseq2 analyses of 6 months old Pitx3(Cre/+);Dot1l(fl/+) mice. (B) Number of genes up/down-regulated with different statistical cut-offs of 6 months old Pitx3(Cre/+);Dot1l(fl/+) mice transcriptomics data. From left to right, Deseq2 adjusted P-values<0.05, or non-adjusted p-values (Wald-tests) >0.01 and <0.05 (C) All retrieved respiratory chain genes at E15.75 sorted neurons. For nuclear-encoded genes, red bars represent (Dot1l(fl/+), gray bars represent Dot1l(fl/fl), and for mtDNA encoded genes: light blue bars represent (Dot1l(fl/+), dark blue bars represent Dot1l(fl/fl). (D) All retrieved mtRibosome encoding genes, red bars represent Dot1l(fl/+), gray bars represent Dot1l(fl/fl). Statistics: ** represent DEseq2, p<0.01 and *p<0.05; ## represent non-adj. P<0.05(Wald-test).

**Supplemental Figure 7 Overview of respiratory genes regulated by Dot1l.**

(A) Respiratory chain complex II-V retrieved from 6 months old Pitx3(Cre/+) induced dissected mouse midbrain transcriptomic profiles. For nuclear-encoded genes, red bars represent (Dot1l(fl/+), gray bars represent Dot1l(fl/fl), and for mtDNA encoded genes: light blue bars represent (Dot1l(fl/+), dark blue bars represent Dot1l(fl/fl). (B) mtRibosome encoding genes retrieved from 6 months old Pitx3(Cre/+) dissected mouse midbrain transcriptomic profiles, red bars represent Dot1l(fl/+), gray bars represent Dot1l(fl/fl). Statistics: ** represent DEseq2, p<0.01 and *p<0.05; ## represent non-adj. P<0.05(Wald-test).

**Supplemental table 1 and 2 Human donor specifications of midbrain material**

