## Supplemental figures and tables for "Neuronal Dot1l is a broad mitochondrial gene-repressor associated with human brain aging via H3K79 hypermethylation"

Supplemental material Heesbeen et al.

Suppl Figure 1

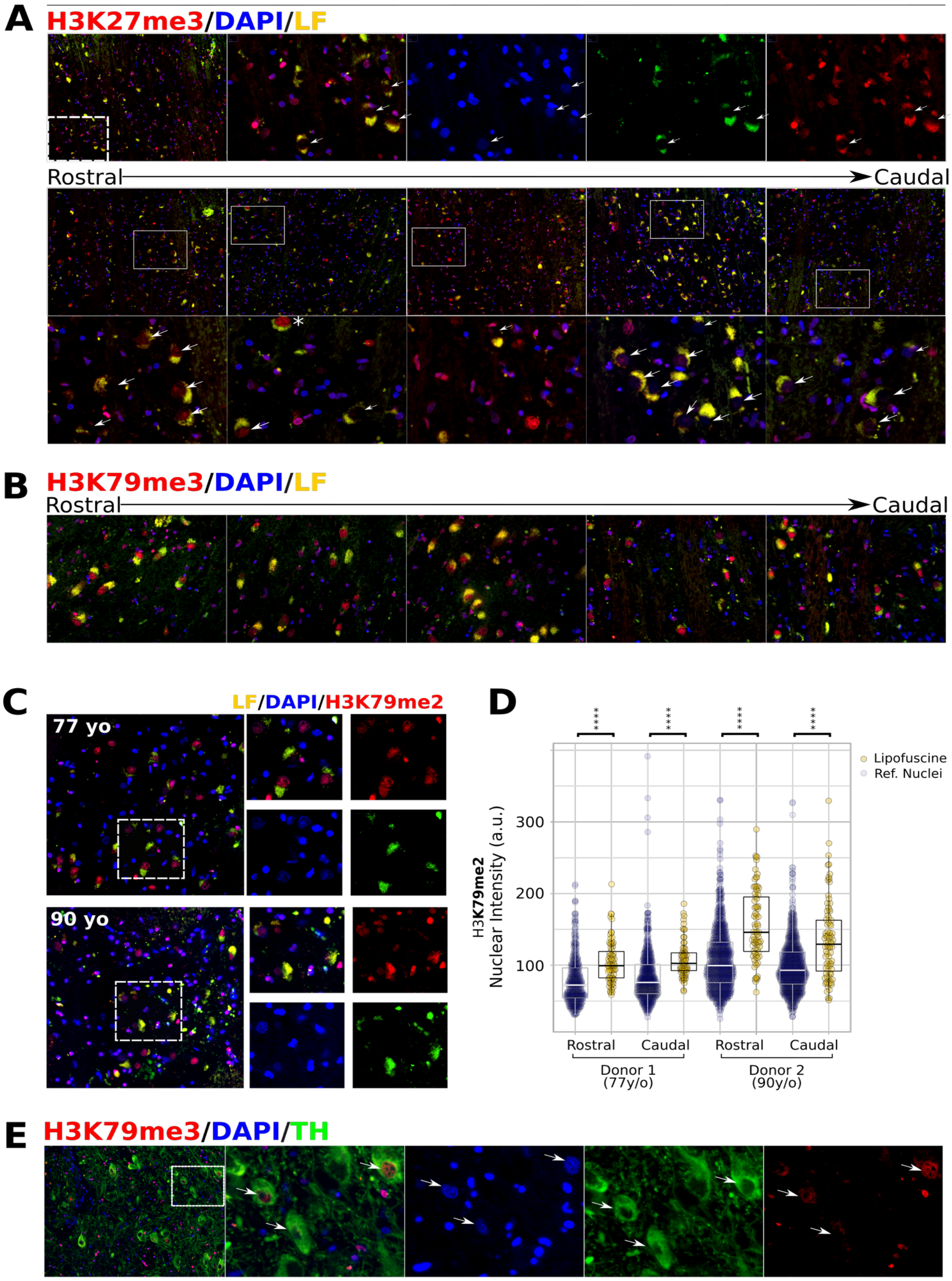

Suppl Figure 2

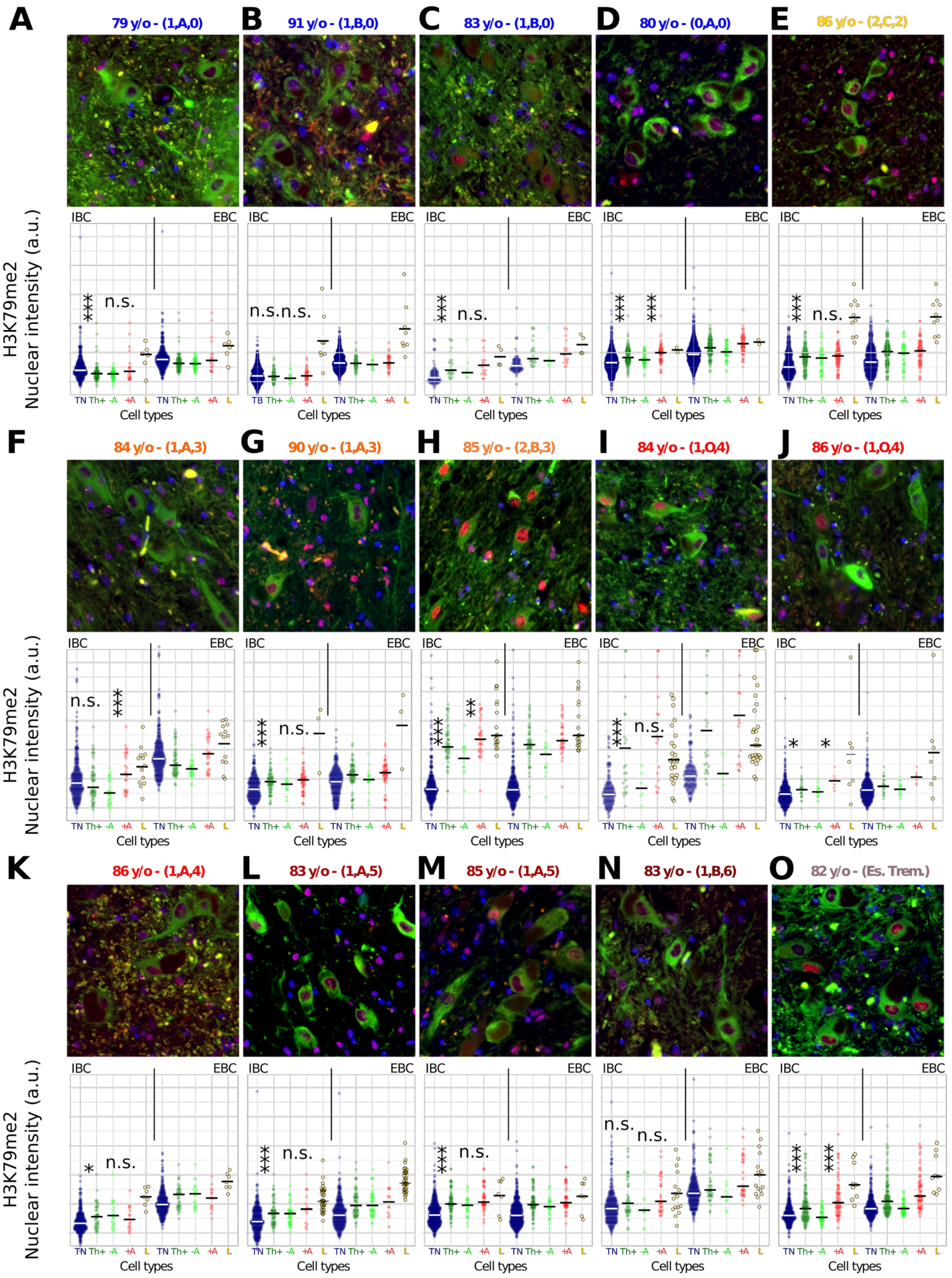

Suppl Figure 3

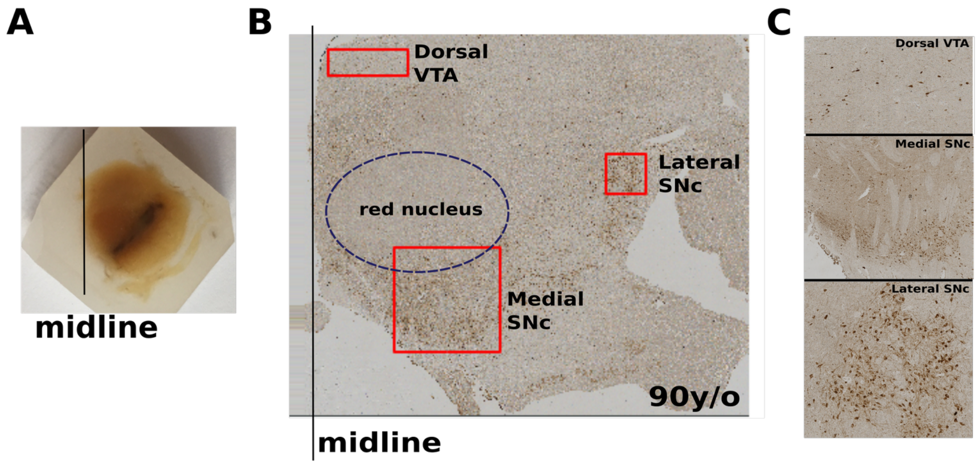

Suppl Figure 4

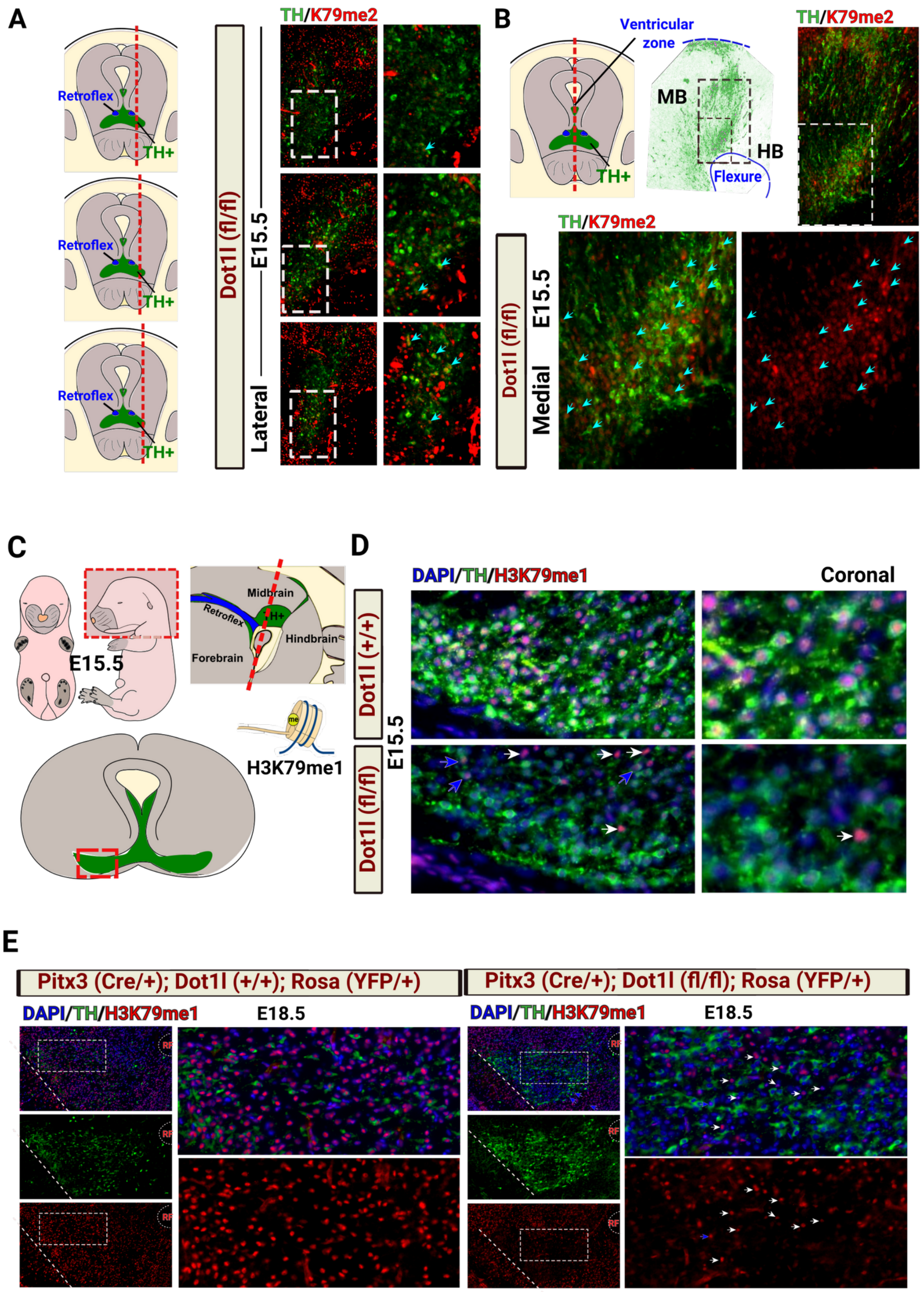

Suppl Figure 5

**A**

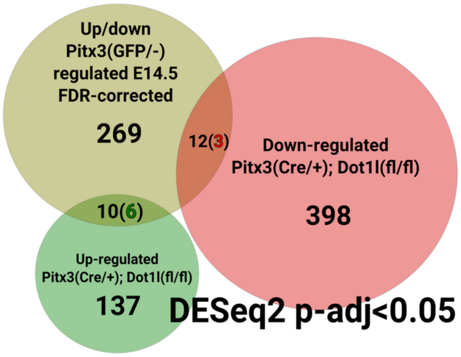


E15.75

**B**

| | RANK PitX3 (GFP/-) | RANK pValue Dot(fl/fl) | Level Dep Rank Dot(fl/+) |
|---|---|---|---|
| Adcy2 | 82 | 153 | No Score |
| Cbln1 | 62 | 514 | No Score |
| Ociad1 | 36 | 302 | No Score |
| Slc32a1 | 16 | 131 | No Score |
| Stxbp6 | 217 | 480 | No Score |
| Tecr | 187 | 518 | No Score |
| **Nr4a2** | **89** | **6** | **191(308)** |
| **VMAT2** | **15** | **192** | **No Score** |
| **Sox4** | **177** | **23** | **306(308)** |

**C**

| UPREGULATED PSEUDOGENES | | Fold Change | | Function related gene (NCBI): |
|---|---|---|---|---|
| 1 | ENSMUSG00000044609 | 1.50591212879062 | **Gm9294** | Ribosomal |
| 2 | ENSMUSG00000081049 | 1.49321433070995 | **Rps24-ps3** | Ribosomal |
| 3 | ENSMUSG00000047905 | 1.46010300850664 | **Gm8566** | Anti-Oxidation(SOD1) |
| 4 | ENSMUSG00000104864 | 1.44869052643099 | **Gm43655** | Unknown |
| 5 | ENSMUSG00000052192 | 1.40316744736745 | **Gm5963** | Unknown |
| 6 | ENSMUSG00000078193 | 1.38975758844883 | **Gm2000** | Ribosomal |
| 7 | ENSMUSG00000067870 | 1.37554894699896 | **Rpl31-ps8** | Ribosomal |
| 8 | ENSMUSG00000104649 | 1.36529028853014 | **Gm43712** | Unknown |
| 9 | ENSMUSG00000056366 | 1.35013250983245 | **Fabp3-ps1** | Fatty Acid Binding |
| 10 | ENSMUSG00000062456 | 1.3493857808012 | **Rpl9-ps6** | Ribosomal |
| 11 | ENSMUSG00000085342 | 1.32561766203794 | **Gm12254** | Unknown |
| 12 | ENSMUSG00000084319 | 1.32419437754443 | **Tpt1-ps3** | tumor protein, translationally-controlled, pseudogene 3 [Source:MGI Symbol;Acc:MGI:2664997] |
| 13 | ENSMUSG00000039617 | 1.31419873945019 | **Gm7488** | RNA processing |
| 14 | ENSMUSG00000084830 | 1.31185163300356 | **Gm14539** | ATP5MD-related |
| 15 | ENSMUSG00000061848 | 1.31102813339557 | **Gm5805** | Long-noncocing |
| 16 | ENSMUSG00000090610 | 1.2810859920404 | **Gm3571** | farnesyl diphosphate synthetase |
| 17 | ENSMUSG00000044751 | 1.27593968016025 | **Gm12231** | ATP synthase, H+ transporting, mitochondrial F0 complex, subunit b, isoform 1 |
| 18 | ENSMUSG00000050299 | 1.27574175998214 | **Gm9843** | Unknown |
| 19 | ENSMUSG00000100863 | 1.27052361052476 | **Gm12669** | glutaredoxin |
| 20 | ENSMUSG00000107383 | 1.2684960909363 | **Gm4366** | Ribosomal (eukaryotic translation elongation factor 1 gamma) |
| 21 | ENSMUSG00000086688 | 1.26733057557459 | **Gm11560** | Ribosomal (YBX1-related) |
| 22 | ENSMUSG00000091549 | 1.25273823646712 | **Gm6548** | Ribosomal (eukaryotic translation elongation factor 1 alpha) |
| 23 | ENSMUSG00000084349 | 1.24435246085593 | **Rpl3-ps1** | Ribosomal |
| 24 | ENSMUSG00000089782 | 1.24021703082236 | **Gm3531** | Ribosomal/transcriptional initiation |
| 25 | ENSMUSG00000101795 | 1.23702244739737 | **Gm5835** | Ribosomal |
| 26 | ENSMUSG00000082691 | 1.22042634424092 | **Dynlt1-ps1** | Dynein |
| 27 | ENSMUSG00000101188 | 1.18738719923614 | **Eif4a-ps4** | Ribosomal (eukaryotic translation elongation factor 1 gamma) |
| **DOWNREGULATED PSEUDOGENES** | | | | |
| 1 | ENSMUSG00000107145 | 0.719633184321127 | Gm43442 | Unknown |

Suppl Figure 6

**A**

| GeneID | RelatedGene |
|---|---|
| 2010107E04Rik | Atp5mpl (mitochondrial) |
| Gm15487 | ATPase, H+ transporting, lysosomal V0 subunit C, pseudogene 2 |
| Gm7308 | Calmodulin 2, pseudogene |
| Gm42418 | Long-Noncoding |
| Gm8129 | ribosomal protein L38 pseudogene |
| Gm10073 | ribosomal protein pseudogene |
| Gm4332 | ribosomal protein S20 pseudogene |
| Gm14303 | ribosomal protein S29 pseudogene |
| Gm14539 | Unknown |
| Gm14586 | Unknown |
| Gm2000 | Unknown |
| Gm5963 | Unknown |

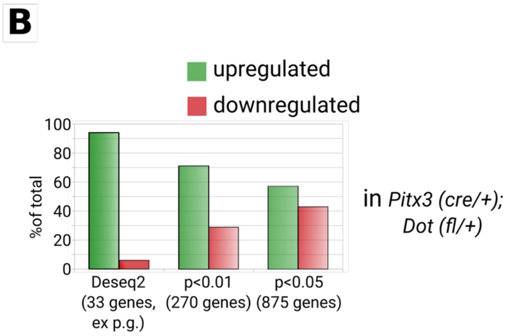


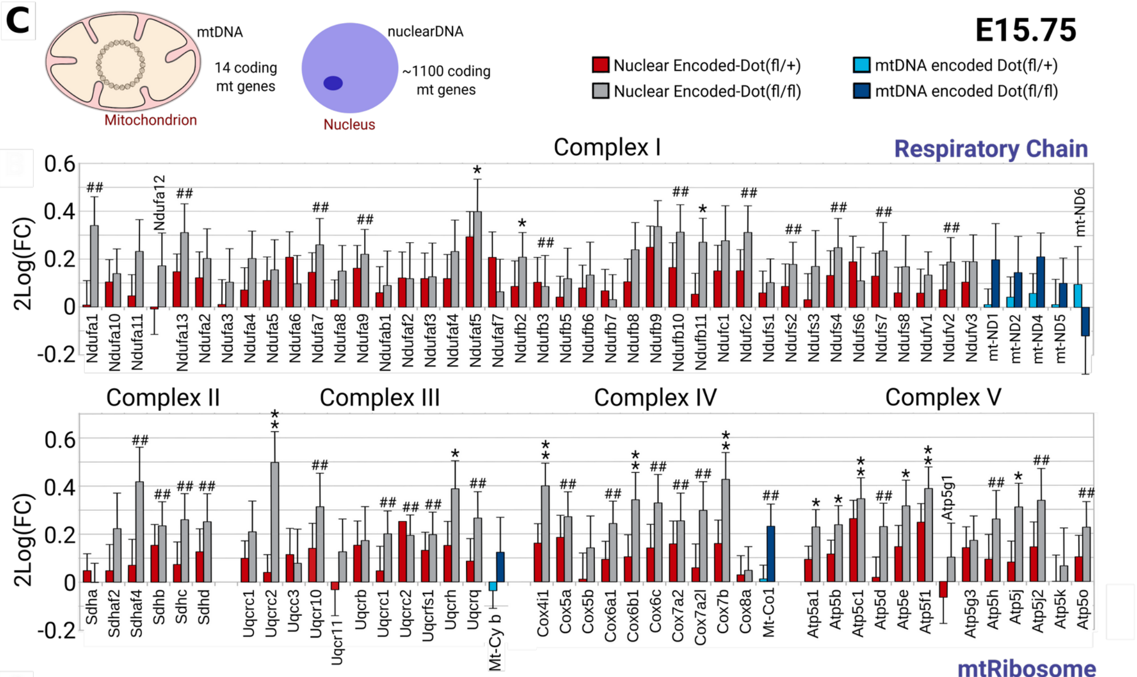


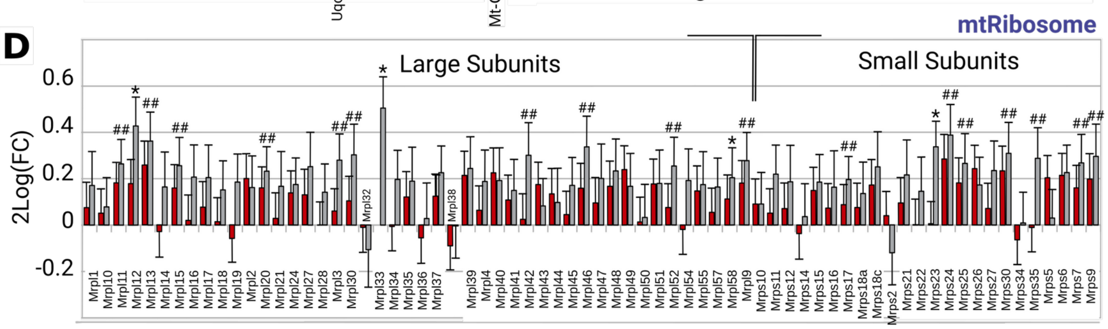

Suppl Figure 7

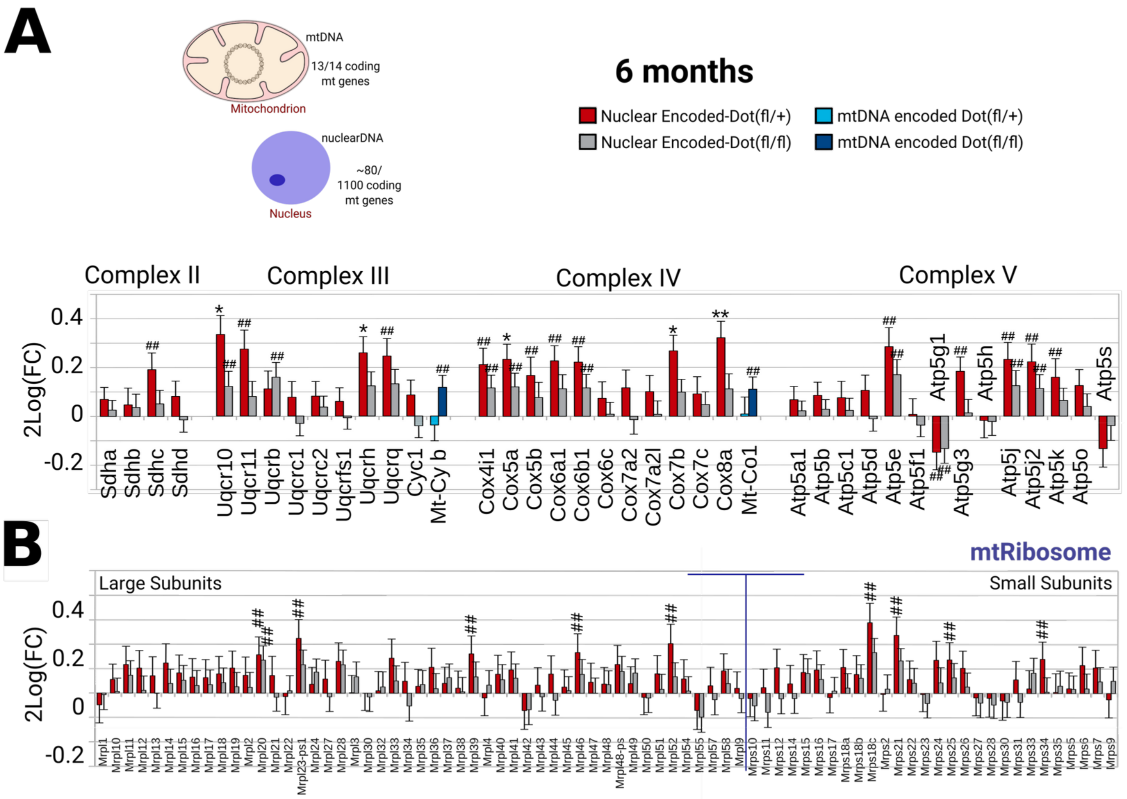

Suppl Table 1 and 2

**Table 1**

| Age | Time of Death | Gender | PMD (hours:min) | pH csf | AD&PD (Tgls, bA, aS) | Region | Cause of Death |
|---|---|---|---|---|---|---|---|
| 45 | | M | 8:50 | 6.71 | 0,0,0 | R1-R2 | |
| 49 | | M | 6:15 | 6.23 | 0,0,0 | R1-R2 | |
| 77 | 09:35 | M | 8:25 | 7.19 | 1,0,0 | R1-R2-C1 | CoD: Perforation of Bladder |
| 90 | 10:20 | F | 6:05 | 6.12 | 3,B,0 | R1-R2-C1 | CoD: Possible infection, Fever unknown foc. |
| 79 | 7:40 | F | 7:40 | 6.02 | 1,A,0 | R2 | CoD:Bronchopneumonia &Sepsis |

**Table 2**

| Age | Time of Death | Gender | PMD (hours:min) | pH csf | AD&PD (Tgls, bA, aS) | Region | Cause of Death |
|---|---|---|---|---|---|---|---|
| 79 | 20:35 | M | 5:20 | 6.72 | 1,A,0 | R1 | CoD:Cachexia |
| 91 | 13:20 | F | 4:15 | 6.5 | 1,B,0 | R2 | CoD: Heart infarction |
| 83 | 22:20 | F | 4:40 | 6.04 | 1,B,0 | R2 | CoD: Ileus with pancreatic cancer |
| 80 | 01:30 | M | 6:30 | 6.43 | 0,A,0 | R2 | CoD: Ventricular fibrillation |
| 86 | 21:00 | F | 6:25 | 6.39 | 2,C,2 | R2 | CoD, Not Known; Diabetis, Breast Cancer, Renal failure |
| 84 | 16:50 | M | 4:50 | 6.41 | 1,A,3 | R2 | CoD: Cachexia/End stage PD |
| 90 | 14:25 | F | 4:50 | 6.37 | 1,B,3 | C1 | CoD Heart Rythm, Resp. Tract Inf. |
| 85 | 10:45 | M | 6:20 | 6.3 | 2,B,3 | R2 | CoD: Cardiac Arrest |
| 84 | 02:40 | M | 6:30 | 6.4 | 1,0,4 | R2 | CoD: Myocard. Inf. Heart fibr. |
| 86 | 02:25 | M | 5:35 | 6.19 | 1,0,4 | R2 | CoD: Aspiration Pneumonia |
| 86 | 14:45 | F | 3:00 | 6.97 | 1,A,4 | R2 | CoD: Seizure by Midazol.&Morf. |
| 83 | 23:15 | M | 6:20 | 6.71 | 1,A,5 | R2 | CoD: CVA (stroke) |
| 85 | 06:59 | M | 8:31 | 6.49 | 1,A,5 | R1 | CoD: Pneumonia&Heartfailure |
| 83 | 01:30 | M | 5:15 | 6.66 | 1,B,6 | R1 | CoD: Pneumonia |
| 82 | 15:07 | F | 7:08 | 6.28 | 1,A,0 | R1 | CoD: Dehydration |
